# Natural language processing charts transcriptome evolution to design combination cancer therapies

**DOI:** 10.1101/2024.04.09.585681

**Authors:** Amir Jassim, Birgit V. Nimmervoll, Sabrina Terranova, Erica Nathan, Katherine E. Masih, Lisa Ruff, Matilde Duarte, Elizabeth Cooper, Linda P. Hu, Gunjan Katyal, Melika Akhbari, Reuben J. Gilbertson, Colton Terhune, Gabriel Balmus, Stephen P. Jackson, Mariella G. Filbin, Anthony Hill, Anarita Patrizi, Neil Dani, Aviv Regev, Maria K. Lehtinen, Richard J. Gilbertson

## Abstract

Combination treatment–the mainstay of cancer therapy–often fails because treatment interactions evoke complex resistance mechanisms that are hard to predict. Designing combination therapy to prevent treatment resistance is especially challenging for rare cancers. Here, we introduce REsistance through COntext DRift (RECODR): a computational pipeline that tracks how genes change their transcriptome context across cancer development and drug treatment conditions. By applying RECODR to a genetically modified mouse model of choroid plexus carcinoma–a rare brain tumour of young children–we identified patterns of transcriptome evolution, cellular heterogeneity and treatment targets that emerged as tumours were initiated and resisted combination treatment. This enabled the prediction of treatment resistance mechanisms and the design of highly effective therapeutic protocols that avoided treatment failure. RECODR can describe complex and dynamic changes in normal and diseased tissues that could be applied to multimodal data from a variety of settings, mitigating treatment resistance across cancers and other diseases.

## INTRODUCTION

Cancers comprise heterogeneous populations of normal and malignant cells that respond differently to treatment, driving complex and dynamic changes in tumour biology that are hard to predict^1,2^. Therefore, selecting curative treatment for the 30-50% of patients who fail first-line therapy is often difficult^3,4^, resulting in high mortality rates among patients with recurrent cancer^5–7^. Different cancers also vary in their dependence on therapeutic targets, further complicating treatment selection: the same mutation can promote or suppress cancer in different tissues and therapies can vary in efficacy among cancers that express the drug target at similar levels^8–10^.

To address these challenges, patients with cancer are usually treated with a combination of different therapies in the hope that this will avoid treatment resistance^11–13^. But combination therapy also often fails, most likely because it does not account for the complex biology that underpins treatment resistance^14^. Designing combination therapy to prevent treatment resistance is especially challenging for rare cancers. The low incidence of these diseases limits patient recruitment to clinical trials and they receive less attention from funders and the research community than more common cancers^15–17^. Consequently, progress in treating rare cancers often stalls, condemning generations of patients and their families to the prospect of unchanging high morbidity and mortality rates. Preclinical models of rare cancers such as genetically modified mouse models can be useful to discover and prioritise new treatments for the clinic^18–24^, but their value for understanding combination treatment and resistance mechanisms is less clear.

The challenges posed by rare cancers are exemplified by childhood brain tumours that kill more children than any other cancer^25^. Choroid plexus carcinoma (CPC) is an especially rare brain tumour of very young children. It accounts for <1% of all brain tumours with an overall age-adjusted incidence rate of 0.008/100,000 people^26,27^. No randomised clinical trials have ever been completed among children with CPC and no new treatments of this cancer have been established in over 50 years^28^. Because CPC is most often diagnosed in very young children, treatment options are further limited by the risk of damage to the developing brain^29^.

Here, we introduce the REsistance through COntext DRift (RECODR) pipeline that detects cancer treatment resistance mechanisms and corresponding drug targets. RECODR deploys graph network (GN) and natural language processing (NLP) analysis of single-cell RNA sequencing (scRNAseq) data to chart how genes change their context within transcriptomes during cancer initiation and treatment resistance (Figure S1). By applying RECODR to chart transcriptome evolution during tumour initiation, monotherapy and combination treatment resistance in a genetically modified mouse model of CPC, we identified treatment targets and resistance mechanisms–including the outgrowth of resistant tumour cell subpopulations–that enabled the design of highly effective therapeutic protocols that prevented treatment failure. Thus, RECODR can describe the complex and dynamic changes in tumour biology that occur during cancer therapy, identifying and mitigating treatment resistance in a manner not possible using conventional genomics, nor feasible through sequential clinical trials.

## RESULTS

### CPC recapitulates choroid plexus lineages

We previously generated a mouse model of CPC that recapitulates the histology, bulk transcriptome, and DNA copy number variations (CNVs) of human CPC and that expresses luciferase, yellow and red-fluorescence proteins (LUC^+^/YFP^+^/RFP^+^; Figure S2A and B)^30,31^. To further characterise the disease, we generated scRNAseq profiles of 2,982 YFP^+^/RFP^+^ CPC cells isolated by stringent fluorescence activated cell sorting (FACS) from allograft tumours in the brains of mice (Figures S2B and C). Thirty-seven percent (n=1,102/2,982) of these profiles closely matched those of progenitor-, immune-, epithelial- or mesenchymal-like cells previously reported among 5,009 scRNAseq profiles of normal mouse embryonic choroid plexus (eCP)^32^–the tissue of origin of mouse CPC^30^ (Figure S2C; Table S1). Thus, like other childhood brain tumours, CPC displays striking similarities to its embryonic tissue of origin^33–36^. Mixed-lineage transcriptomes were observed in the remaining 1,880 CPC cells, compatible with aberrant tumour cell differentiation. CPC scRNAseq profiles also inferred the presence of multiple CNVs and intrachromosomal translocations that we validated in FACS-isolated YFP^+^/RFP^+^ CPC cells using spectral karyotyping and in intact tumours using fluorescence *in situ* hybridization (Figures S2D to H).

### The RECODR pipeline

Differential expression analysis (DEA), which determines which genes are expressed at different levels between conditions, can be used to define cell types and their associated functions. But DEA has limited power to decipher patterns of gene expression among cells with similar transcriptomes– such as those that constitute the bulk of CPC (Figure S2C)–or to describe complex changes in gene expression relationships that occur during differentiation, malignant change and treatment resistance. More sophisticated computational approaches to identify drug targets from large datasets have been developed; but these compare gene expression in test tissues to canonical pathways and interactomes defined previously in other contexts that may not be applicable to the disease of interest^37^.

To overcome these challenges, we developed the RECODR pipeline that integrates GNs and NLP to chart transcriptome evolution in scRNAseq profiles during tumour development and treatment resistance, predicting patterns of cell plasticity and pinpointing therapies to mitigate resistance (Figure S1; STAR methods). Rather than organizing cells in two-dimensional space according to their overall levels of gene expression, RECODR organizes genes through GNs in two-dimensional space according to their pattern of co-expression across all cells in the tissue. Genes with tightly correlated expression across cells, and therefore potentially related function, are clustered closely together forming communities. RECODR then deploys Node2Vec^38^ to take biased random walks across these GNs to sample the transcriptome context of each gene. This generates an input format that recapitulates the structure of each GN that is then fed into the skip-gram model of Word2Vec^39^. Word2Vec is a neural network designed to learn the context of words in large bodies of text that RECODR uses to learn the context of each gene within each GN, providing a rich semantic view of the entire tissue transcriptome. By comparing the context of each gene between GNs, RECODR identifies vectors with high cosine similarities that preserve their transcriptome context and those with lower cosine similarities whose context has drifted. Our underlying hypothesis is that genes displaying a significant shift in transcriptome context during treatment resistance, regardless of what is known about that shift from prior databases or biological contexts, have undergone a significant drift in biological function and may therefore be a drug target capable of mitigating resistance. This approach has the advantage of predicting drug targets based on the gene context within the studied transcriptome, rather than fitting potential targets to gene expression patterns previously defined in unrelated contexts. By deploying this and other graph-based metrics, RECODR generates a ‘target score’ that prioritises each gene as a potential drug target to mitigate resistance.

### Transcriptome evolution during CPC tumorigenesis

To understand changes in the biology of eCP as it transformed into CPC, we analysed the scRNAseq profiles of these tissues using RECODR. In keeping with its ontogeny, 91% (n=2,031/2,235) of the genes present in the RECODR-generated GN of CPC (R^CPC^-GN) were also present in the R^eCP^-GN (n=9,389; representation factor for overlap, p=2.4e-243; Figures S3A and B; Tables S2 to S4). However, the R^CPC^-GN was reduced markedly in size and displayed significant differences in community structure relative to the normal parent tissue, indicating that gene-co-expression patterns differed significantly between these two tissues (Figure 1A to C). For example, RECODR replaced a large community of genes associated with mature ciliated epithelium in the R^eCP^-GN with a smaller community of immature choroid plexus/CPC-associated genes in the R^CPC^-GN that was populated by genes from all R^eCP^-GN communities. Following transformation, RECODR also switched the community-type location of 41% (n=469/1,197) of genes present in communities enriched for genes associated with progenitor, immune, or mesenchymal cells. These genes were assigned relatively low NLP cosine scores indicating an altered transcriptome context. They enriched the R^CPC^-GN progenitor community with genes associated with RNA translation–a key property of neural progenitor cells^40,41^–and the immune community with regulators of innate immunity and microglial cell function (Figure 1C to F; Table S4). The 728 genes that remained in the same communities following transformation displayed relatively high NLP cosine scores indicating a retained transcriptome context, suggesting they supported similar functions in the normal parent and daughter malignant tissue (Figure 1C to I). Sixty percent (n=435/728) of these genes enriched the progenitor communities of the R^eCP^- and R^CPC^-GNs with regulators of cell proliferation, nucleotide excision repair, and DNA double strand break repair (Figure 1C and G; Table S4).

**Figure 1.**
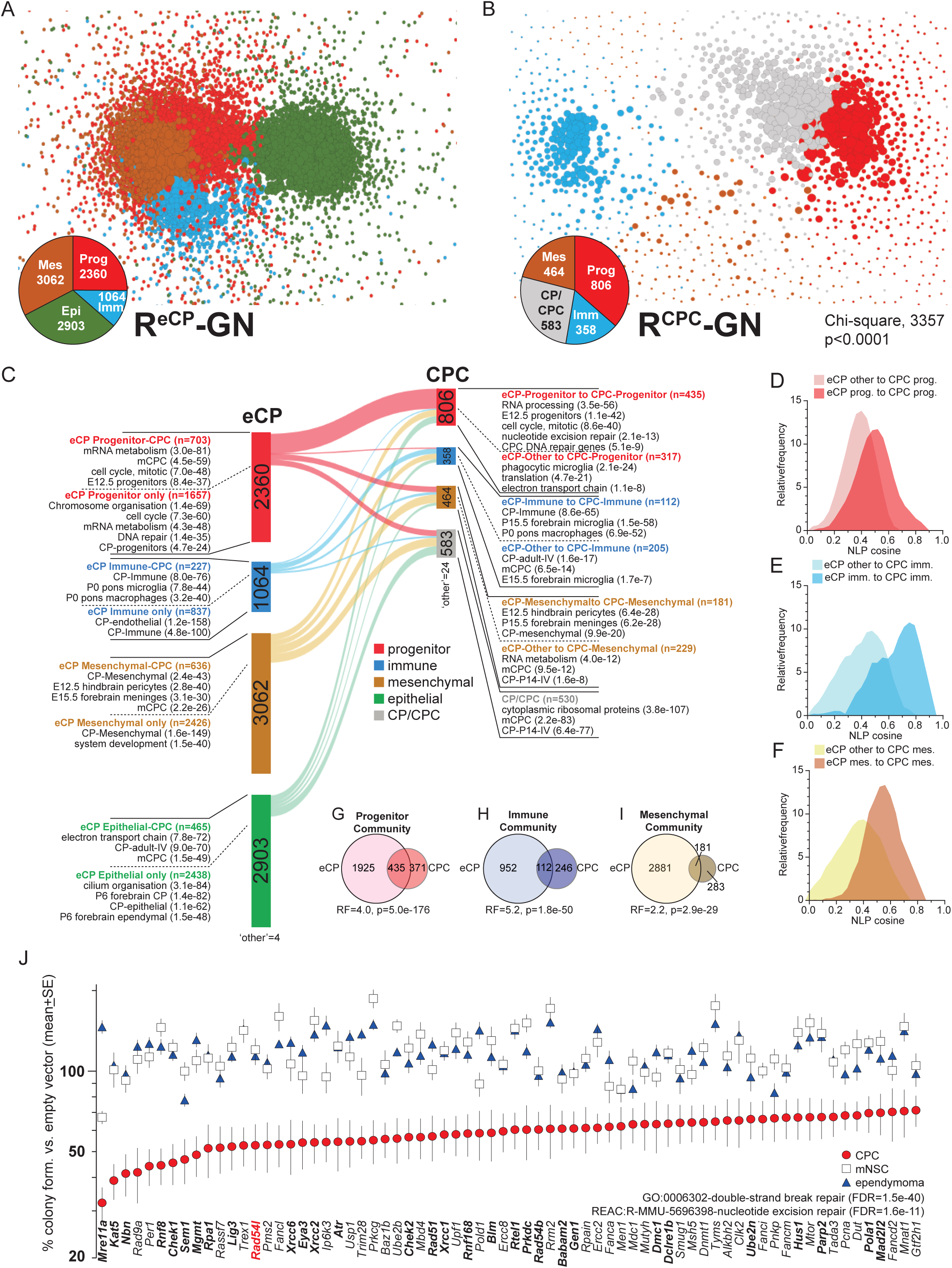
RECODR reveals DNA repair as a CPC vulnerability. RECODR generated graph networks (GNs) of scRNAseq profiles of **(A) e**CP and **(B)** untreated CPC. Pie charts indicate the number of genes in progenitor (Prog), immune (Imm), mesenchymal (Mes), epithelial (Epi) or CP/CPC communities. Chi-square reports differences in gene numbers in communities between (GNs). **C**. Alluvial plot of gene locations between communities in the R^eCP^- and R^CPC^-GN. Enriched pathways in relevant gene groups are shown with FDR (Tables S2 and S3). Frequency plots of NLP cosine scores reporting transcriptome shifts of the indicated gene groups in R^CPC^-GN progenitor **(D),** immune **(E),** and mesenchymal **(E)** communities. Venn diagrams of overlap in genes in progenitor **(G),** immune **(H),** and mesenchymal **(I)** communities of R^eCP^- and R^CPC^-GNs with representation factor (RF) and associated p-value for overlap progenitor. **J**. Graph of percent colony formation by CPC, mouse neural stem cells (mNSCs) and mouse *RTBDN-*driven ependymoma following shRNA silencing of the indicated DNA repair transcript relative to matched-control transduced cells. Only genes demonstrating significant (p<0.05) growth suppression in CPC cells are shown. Enriched pathways in CPC dependency-genes are shown.

To validate our findings, we performed RECODR analysis of single nuclear RNAseq profiles of human CPC which produced a very similar GN community structure to the mouse R^CPC^-GN; confirming that our model faithfully recapitulates key elements of the human disease, and that our pipeline identifies similar biological processes across species (Figure S3C; Tables S5).

As expected, RECODR-GN communities were highly enriched in scRNAseq profiles expressing the corresponding and previously reported^32^ eCP progenitor-like, immune-like, or mesenchymal-like signatures (compare Figures S2C and S3D; Table S6). However, DEA among tumour cells defined using these previously published eCP signatures detected only a handful of differentially expressed genes and provided little insight into gene co-expression patterns (Figure S3E; Table S7). Thus, compared to DEA, RECODR appears to provide a much richer view of changing gene expression relationships during transformation.

Together, these data support the notion that CPCs arise following restricted and aberrant differentiation of eCP, generating tumours that lack mature ciliated epithelial cells–a feature supported by our previous histological studies^30^. Progenitor-like cells in normal eCP have previously been shown to generate diverse cell types including epithelia and neurons^32^; therefore, CPC cells enriched for progenitor community expression are likely the apex of the malignant cell hierarchy.

### DNA repair is a therapeutic vulnerability of CPC

The prominence of DNA repair genes identified within the R^eCP^- and R^CPC^-GN progenitor communities, and the high frequency of inter-chromosomal translocations in CPC (Figure S2E to G)– an established feature of disordered double-strand break repair–suggested that DNA repair might be a therapeutic vulnerability of CPC. Indeed, we showed previously that *Taf12* and *Rad54l* that regulate nucleotide excision^42^ and homologous recombination repair^43^, respectively, are required for CPC formation *in vivo*^30^.

As a first step to test if DNA repair is a therapeutic vulnerability of CPC, we measured CPC cell clonogenic colony formation *in vitro* following knock down of 227 DNA repair genes. Thirty-one percent (n=70/227) of these genes were required for CPC cell colony formation and were highly enriched for mediators of nucleotide excision and double-strand break and repair, including *Rad54l* (Figure 1J; Table S8 and S9). Conversely, only one (*Mre11a*) and four (*Sem1, Pnkp, Men1, Mdc1*) genes were required for colony formation by mouse neural stem cells and ependymoma cells^44^, respectively.

To test more directly if DNA repair is a therapeutic vulnerability of CPC, we treated mice with these tumours with four different regimens of the blood brain barrier penetrating, Ataxia-Telangiectasia Mutated (ATM) inhibitor, AZD1390^45,46^. We compared this therapy to various doses and schedules of image-guided, fractionated radiotherapy that is used to treat older children and adults with CPC in the clinic (Figure S4A to C). Remarkably, AZD1390 monotherapy significantly suppressed CPC growth, supporting the notion that the disease is peculiarly dependent on DNA repair. To our knowledge, this is the first report of single agent ATM inhibitor efficacy in cancer. Like human CPC, the mouse disease was also sensitive to radiation therapy.

### Transcriptome evolution during monotherapy resistance

Despite an initial therapeutic response to AZD1390 or radiation, all mice ultimately relapsed and succumbed to their disease (Figure S4B and C). To begin to understand how CPCs resisted these treatments, we used RECODR to compare transcriptomic changes among scRNAseq profiles of CPC cells that we FACS-isolated from tumours that had relapsed during regimen AZ^1390^-4 (n=48,068 cells) or irradiation regimen IR-2 (n=17,544 cells) as well as those of eCP and control CPC.

Despite the different modes of action of AZD1390 and radiation, RECODR generated strikingly similar R^AZD1390^- and R^IR^-GNs (Figure 2A and B; Table S10 and S11). These GNs differed significantly from the R^CPC^-GN, suggesting a common mechanism(s) underpinned resistance to both treatments. Notably, the progenitor communities in both the R^AZD1390^-GN (62%, n=2,509/4,035 of GN genes) and R^IR^-GN (59%, n=2,055/3,509) were markedly expanded relative to the R^CPC^-GN (36%, n=806/2,235; Chi-square, p<1e-15 for both comparisons; Figure 2A to C; Table S10 and S11). The genes in these expanded progenitor communities were derived from four main sources. First, the great majority of genes within the R^CPC^-GN progenitor community (n=806) were also placed in the R^AZD1390^- (n=748 genes) and R^IR^-GN (n=694 genes) progenitor communities: more than half of these genes were also present in the R^eCP^-GN progenitor community. RECODR assigned high NLP cosine scores to these genes, suggesting that their associated functions–including RNA processing, mitosis and DNA repair–are retained, core properties of progenitor-like cells in eCP, untreated CPC, and relapsed tumours (Figures 2C to F; Table S12). RECODR also placed genes from other R^CPC^-GN communities within the progenitor communities of relapsed CPC. These included almost all CP/CPC community genes and around one-half of those in the mesenchymal community; enriching R^AZD1390^- and R^IR^-GN progenitor communities with ribosomal protein and electron transport chain regulators.

**Figure 2.**
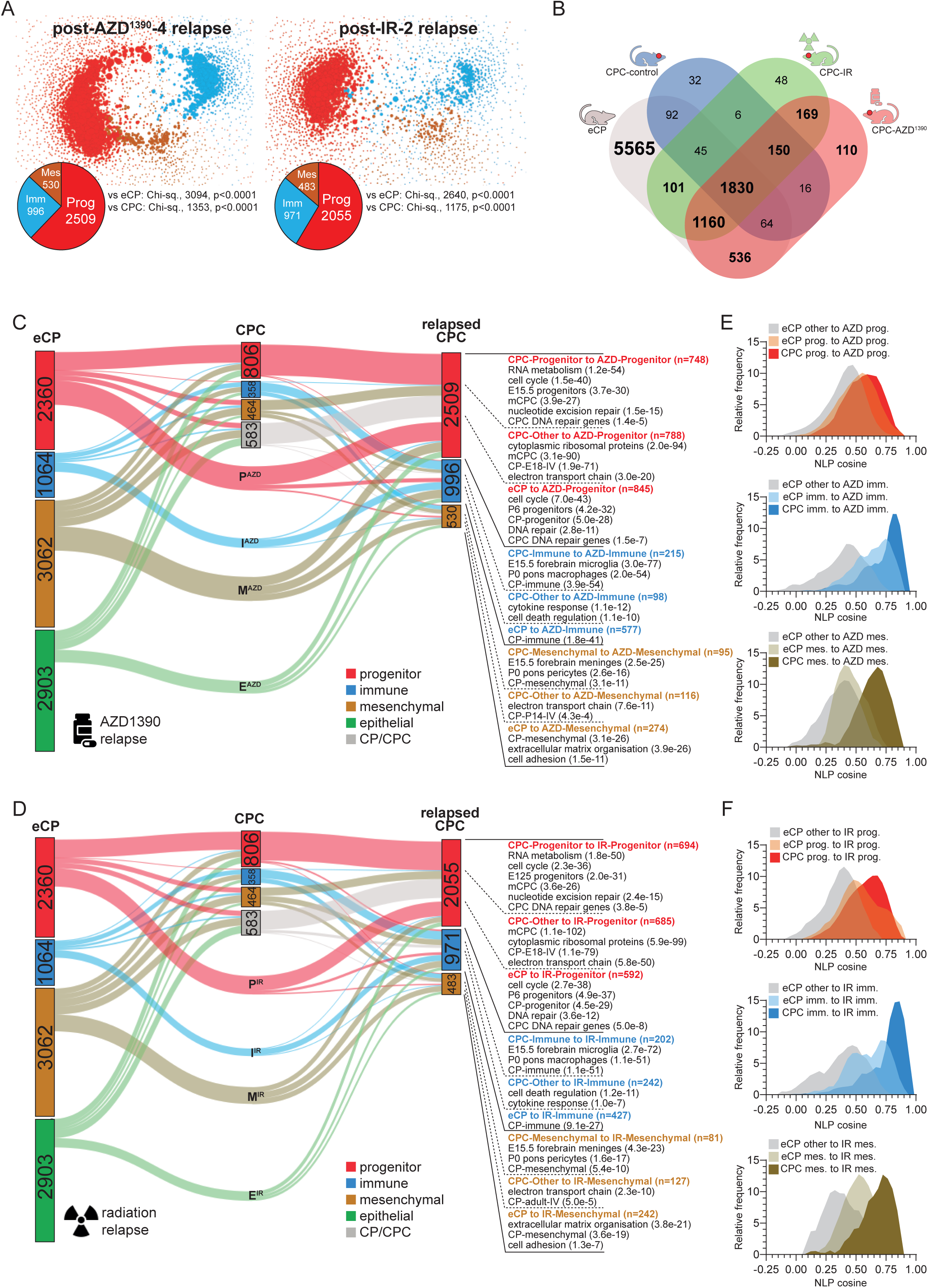
RECODR reveals CPC transcriptome evolution following monotherapy resistance. **A.** RECODR generated graph networks (GNs) of scRNAseq profiles of CPC that relapsed during AZD1390 (left) or irradiation (right) therapy. **B.** Chi-square reports differences in gene numbers in communities between relapsed CPC and control CPC or eCP GNs. Venn diagram of GN genes in the indicated GNs. Alluvial plots of gene locations between communities in **(C)** R^eCP^-, R^CPC^, and R^AZD1390^-GN, and **(D)** R^eCP^-, R^CPC^-, and R^IR^-GN. Enriched pathways in relevant gene groups are shown with FDR (Tables S10 to S12). Frequency plots of NLP cosine scores reporting transcriptome shifts of the indicated gene groups in **(E)** R^AZD1390^-GN and **(F)** R^IR^-GN progenitor, immune, and mesenchymal communities.

Remarkably, 845 and 592 genes placed by RECODR within the R^AZD1390^- and R^IR^-GN progenitor communities, respectively, were not present in the R^CPC^-GN; but these genes were located on R^eCP^- GN–the majority within its progenitor community (Figure 2C and D; Table S12). These genes had high NLP cosine scores, supporting the notion that resistance of CPC to AZD1390 or radiation included the reacquisition of functions operating in the ancestral, normal embryonic tissue (Figures 2E and F; Tables S10 to S12). These genes provided the progenitor communities of relapsed CPC with additional members of the cell cycle e.g., *Cdk6*, *Cdk7*, *Ccna2*, *Ccnb2*, and *Ccnd3* (Cell Cycle, REAC:R-MMU-1640170: adjusted p=4.0e^-48^) and DNA repair machinery e.g., *Parp1*, *Chek1*, *Rad50*, *Rad51*, *Rad21* and *Mre11* (DNA repair, GO:0006281: adjusted p=1.1e^-13^). Finally, RECODR placed a small number of unique genes in the AZD^1390^- and IR-GNs although these were not enriched for any clear function (Table S10 and S11).

Mapping RECODR expression enrichment scores of GN communities to individual CPC scRNAseq profiles confirmed that progenitor community expansion in the R^AZD1390^- and R^IR^-GNs was driven by a significant and selective increase in the proportion of progenitor-like cells (Figures S5A and B; Tables S13 and S14). Indeed, expression of the protein product of *Dhfr*–a progenitor community gene that regulates human and mouse neural progenitors^47^–was increased significantly in relapsed GFP^+^ CPC cells, while nuclear expression of γH2AX that marks DNA double strand breaks was decreased (Figure S5C to F).

Thus, RECODR predicted that CPCs resisted AZD1390 and radiation through a common mechanism associated with the outgrowth of progenitor-like tumour cells possessing increased proliferative and DNA repair capacity, obtained in part, through the reacquisition of functions active in the ancestral, normal tissue. RECODR detected much smaller, non-significant changes in the immune and mesenchymal communities of both the R^AZD1390^- and R^IR^-GNs, and immune- and mesenchymal-like cells were not enriched in relapsed CPCs relative to controls (Figure S5A and B).

### RECODR predicts *PARP1* as a shared monotherapy resistance target

To determine which genes might serve as therapeutic targets to mitigate monotherapy resistance, we reviewed the RECODR ‘target scores’ that ranks the context drift of genes between GNs to detect potential mediators of treatment resistance (STAR methods). *Parp1,* a relapsed progenitor community gene that was ‘reacquired’ from the R^eCP^-GN but absent from the R^CPC^-GN, ranked in the top 5% of both AZD1390 and radiation resistance targets (Figures 3A and B; Tables S15 and S16). *Parp1* scored particularly highly as a radiation resistance target since its connectome (partners within two hops) reached 63% (n=1,310/2,055) of all genes in the R^IR^-GN progenitor community (Figure 3C and D). In stark contrast, DEA ranked *Parp1* in the bottom 50% of differentially expressed genes in AZD1390 and radiation-resistant progenitor-like cells (0.39 and 0.43 log2 fold difference respectively) relative to progenitor cells in untreated CPC (Figure 3E and F; Tables S17 and S18).

**Figure 3.**
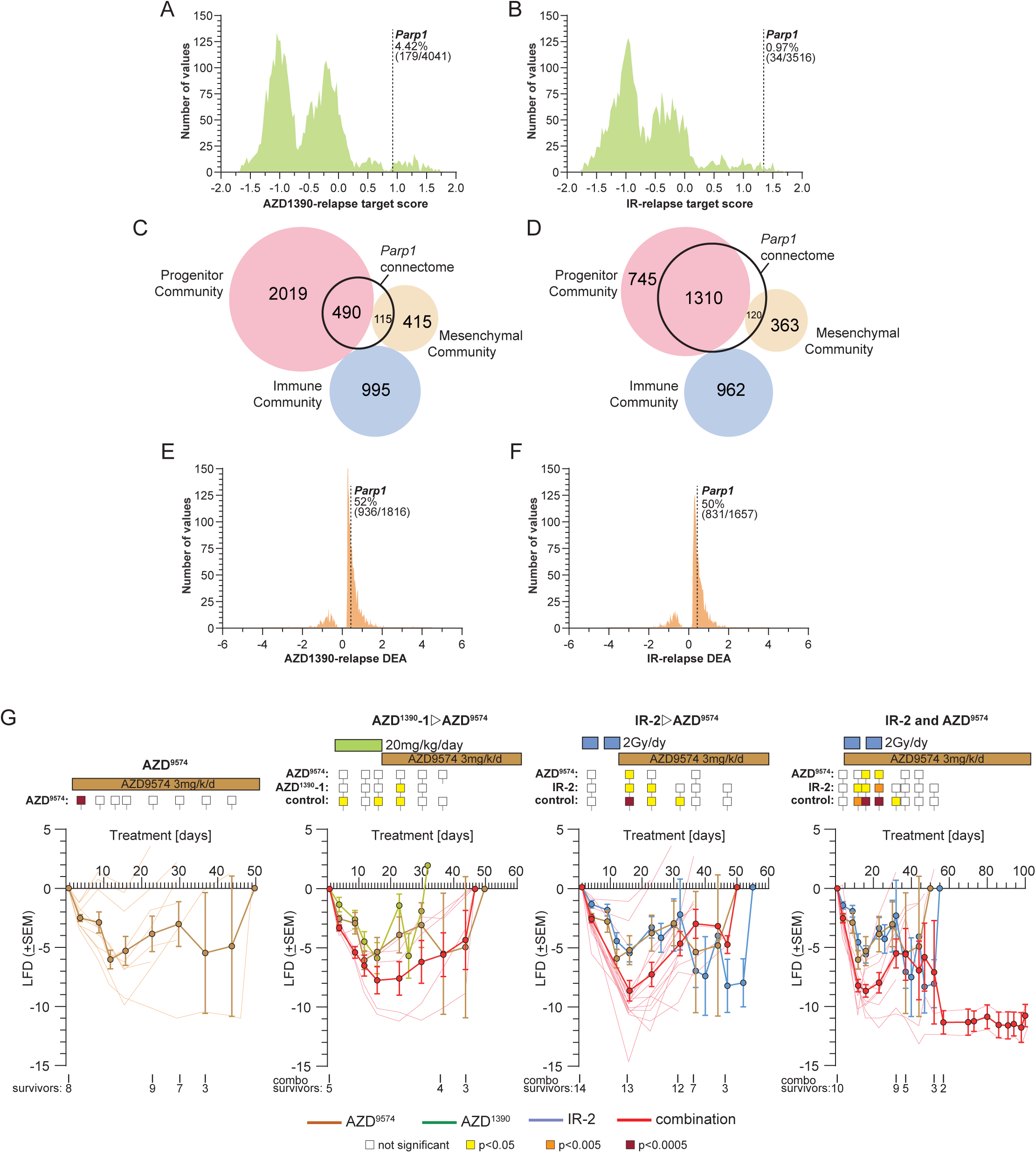
RECODR identification of drug targets in monotherapy relapsed tumours. Frequency plots of ‘target scores’ of genes on the **(A)** R^AZD1390^-GN and **(B)** R^IR^-GN. The percentage ranking of *Parp1* is shown in each. Venn diagram of overlap *Parp1* with communities in the **(C)** R^AZD1390^-GN and **(D)** R^IR^-GN. Frequency plots of significantly differentially expressed (FDR<0.05) genes between progenitor-like cells in **(E)** AZD1390 relapsed and **(F)** radiation relapsed CPC versus progenitor cell in control CPC. The percentage ranking of Parp1 is shown in each. **G.** (Top) Different treatment protocols of CPC in mice with the PARP1 inhibitor alone or with AZD1390 treatment or irradiation treatment. Coloured squares report significance (Wilcoxon adjusted p-value) of treatment-induced growth suppression relative to controls in plots below. (Bottom) Graphs of the average log fold difference (±standard error of mean) of each treatment relative to control treated CPC. Numbers of surviving mice at the indicated times for combination (combo) treatment are shown below each graph.

To test directly if PARP1 contributed to AZD1390 and/or radiation resistance of CPC, we treated mice with these tumours with the blood brain barrier penetrating PARP1 inhibitor AZD9574 either alone or following an initial response to AZD1390 or radiation (Figures 3G). The addition of AZD9574 treatment produced significantly greater and more prolonged periods of tumour growth suppression than either monotherapy alone, although these responses were short-lived (Figure 3G). Toxicity precluded concurrent dosing of AZD9574 and AZD1390; however, the combination of AZD9574 with radiation produced even greater and more sustained levels of tumour growth suppression, with 20% (n=2/10) of mice achieving complete remission of their disease (Figure 3G). These observations provided initial proof-of-principle that RECODR might be useful for pinpointing treatment resistance mechanisms.

### Transcriptome evolution during combination therapy resistance

Understanding how tumours resist single therapies may have value, but cancers are almost always treated with combinations of different treatments. Interaction among these treatments drives both efficacy and resistance mechanisms that can be hard to predict, undermining the design of effective treatment protocols. Therefore, we looked to see if RECODR might help design more effective treatment combinations.

First, we identified the most effective combination of AZD1390 and radiation to treat CPC in our mice: this combination is being tested in adult patients with glioblastoma (NCT03423628). Sixty-five mice with CPC were randomized among seven different combinations of AZ1390 and radiation, ranging from low to high treatment intensity (Regimens A-G; Figure 4). Protocols involving at least 2Gy/day/daily for 5 days of radiation together with 20mg/kg/day of AZD1390 markedly inhibited tumour growth. Among these, Regimen-E was the most effective at suppressing tumour growth relative to control, AZD1390 or radiation monotherapy; however, all mice eventually relapsed and succumbed to their disease.

**Figure 4.**
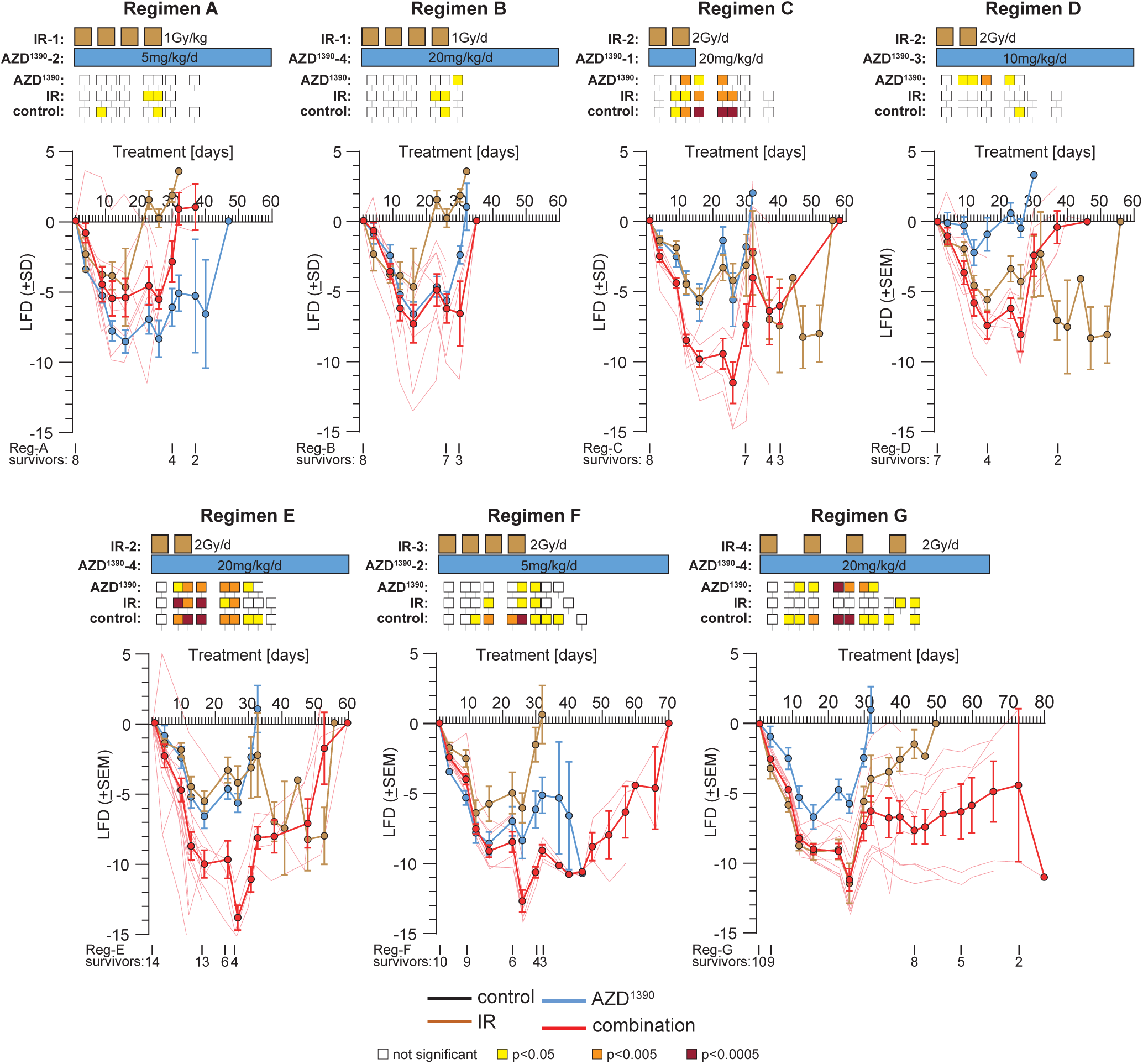
Combination AZD1390 and irradiation treatment regimens. (Top in each) Different combination treatment protocols of CPC in mice with AZD130 and irradiation. Coloured squares report significance (Wilcoxon adjusted p-value) of treatment-induced growth suppression relative to controls in plots below. (Bottom) Graphs of the average log fold difference (±standard error of the mean) of each treatment relative to untreated CPC. Numbers of surviving mice at the indicated times for combination treatments are shown below each graph.

To understand how mice resisted the combination therapy, we used RECODR to compare transcriptomic changes among 8,032 scRNAseq profiles of CPC cells that we FACS-isolated from mice that relapsed during Regimen-E treatment (RegE-GN), with those of eCP, control CPC, and monotherapy relapsed CPC. Despite the common pattern of transcriptome evolution of CPCs resisting AZD1390 and radiation monotherapies, RECODR generated a R^RegE^-GN that differed both in size and structure from all GNs, suggesting that the combination therapy evoked a different resistance mechanism than that driven by these treatments delivered as monotherapies (Figure 5A and B; Table S19). Similar to the R^AZD1390^- and R^IR^-GNs, the progenitor community of the R^RegE^-GN was significantly expanded relative to that of the R^CPC^-GN and comprised 48% (n=2,909/5,849) of all genes in the GN. Eighty-one percent (n=2,344/2,909) of genes placed by RECODR in the R^RegE^-GN progenitor community, were common to the progenitor communities of the R^AZD1390^- and/or R^IR^-GN: these included a ‘core group’ of 391 genes present in the progenitor communities of all GNs (Figures 5B and C and S6A; Tables S12 and S19). RECODR assigned these genes high NLP cosine scores, and they enriched all progenitor communities, including that in the R^RegE^-GN, with regulators of RNA translation, cell cycle and DNA repair (Figures 5C and D; Table S12). Thus, RECODR identified these as core functions of eCP and its malignant daughter tumours, both before and following treatment relapse. The proportion of tumour cells enriched for expression of progenitor community gene transcripts and DHFR protein was also significantly increased in Regimen-E relapsed tumours, while γH2AX nuclear expression was decreased (Figures 5E and F, S5C to F). Thus, AZD1390 and radiation, whether delivered alone or in combination, drives the outgrowth of progenitor-like tumour cells that likely contribute to CPC resistance.

**Figure 5.**
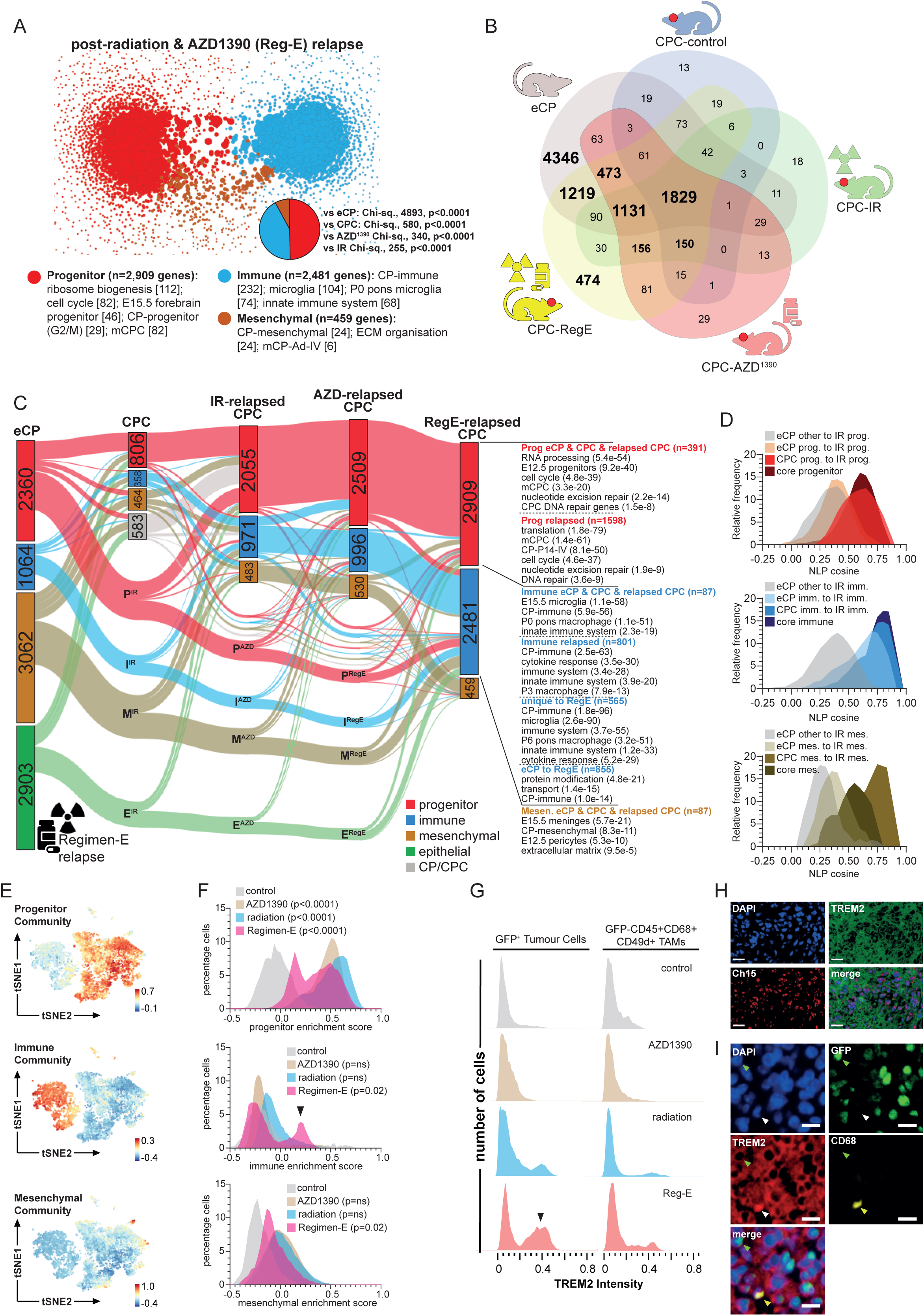
RECODR reveals CPC transcriptome evolution and outgrowth of immune-like CPC cells during Regimen-E combination treatment resistance. **A.** RECODR generated graph networks (GN) of scRNAseq profiles of CPC that relapsed during Regimen-E treatment. Chi-square reports differences in gene numbers in communities between Regimen-E relapsed CPC and the indicated GNs. **B.** Venn diagram of GN genes in the indicated GNs. **C.** Alluvial plot of gene locations between communities in R^eCP^-, R^CPC^-, R^IR^-, R^AZD1390^-, and R^RegE^-GNs. Enriched pathways in relevant gene groups are shown with FDR (Tables S12 and S19). **D.** Frequency plots of NLP cosine scores reporting transcriptome shifts of the indicated gene groups. **E.** t-distributed Stochastic Neighbor Embedding (tSNE) plots of the indicated community gene signature enrichment score in scRNAseq profiles of CPC cells during relapse to Regimen-E. **F.** Frequency plots of community enrichment scores in scRNA seq profiles across the indicated conditions. Arrow indicates subpopulation of immune community highly enriched cells. **G.** Frequency plots of TREM2 immunofluorescence across different conditions and control (untreated CPC) in GFP^+^ CPC cells or GFP^-^/CD45^+^/CD69^+^/CD49^+^ tumour associated macrophages (TAMs). Arrow indicates subpopulation of TREM2 high expressing CPC cells. **H.** Representative concurrent TREM2 immunofluorescence and chromosome 15 FISH of Regimen-E treated CPC. Scale bar=20µm. **I.** Co-immunofluorescence of GFP, TREM2 and CD68 in Regimen-E relapsed CPC showing GFP^+^ CPC cells and GFP^-^/CD68^+^ TAM. Scale bar=20µm.

But RECODR also identified a dramatic increase in the size of the immune community in Regimen- E relapsed CPC that was not seen when either AZD1390 or radiation were administered alone. RECODR placed 42% (n=2,481/5,849) of all R^RegE^-GN genes in the immune community, compared to only 25% (n=996/4,035; Chi-square p<1e^-15^), 28% (n=971/3,509; Chi-square p<1e^-15^) and 16% (n=358/2,211; Chi-square p<1e^-15^) of genes in the immune communities of the R^AZD1390^-, R^IR^- and R^CPC^-GNs, respectively (Figures 5A to C and S6B; Table S19). This expanded R^RegE^-GN immune community included the majority of R^CPC^-GN (n=258/358), R^AZD1390^-GN (n=926/996) and R^IR^-GN (n=701/971) immune community genes, enriching the immune communities in each of these GNs with regulators of microglia, macrophages, and the innate immune response, suggesting these are core properties of CPC, whether untreated or relapsed (Figure 5C and D and Figure S6B; Table S19). Remarkably, 50% (n=1,244/2,481) of R^RegE^-GN immune community genes were not placed by RECODR in any other tumour GN: 69% (n=855/1,244) of these genes were ‘reacquired’ from the R^eCP^-GN, while the remaining 389 genes were unique to the R^RegE^-GN. These groups of genes further enriched the R^RegE^-GN immune community with regulators of the innate immune system, including microglia and macrophages. Mapping RECODR expression enrichment scores of GN communities to individual CPC scRNAseq profiles confirmed that immune community expansion in the R^RegE^-GN was driven by a significant and selective increase in a subpopulation of cells defined by high expression of immune community genes (Figures 5E and F; Table S20).

Given that CPC scRNAseq profiles were generated from FACS-isolated tumour cells, then we reasoned that these immune-like cells may be tumour cells expressing a myeloid mimicry programme, similar to that previously reported in glioma^48^. Indeed, RECODR detected enrichment of a glioma myeloid mimicry signature in the R^RegE^-GN immune community (MESImm_Gene_Signature, p=1.8e-09; Table S19). However, it remained possible that inadequate separation of CPC cells from brain resident microglia and/or tumour associated macrophages contributed to this immune-like subpopulation. Indeed, *Triggering receptor expressed on myeloid cells 2* (*Trem2*) that marks microglia, glioma associated macrophages, and certain cancer cells^49–51^ was expressed in all CPC cells enriched for this community type (Figure S6D).

To distinguish these possibilities, we conducted extensive, multiplex profiling of cell surface antigens and CNVs across all tumour states to identify tumour cells (TREM2^+^/GFP^+^/Chromosome 15 FISH gain), tumour-associated macrophages (GFP^-^/TREM2^+^/CD45^+^/CD68/P2Y12^+^/CD49d^+^) and microglia (GFP^-^/TREM2^+^/CD45+/CD68/P2Y12^+^/CD49d^-^; Figure 5G to I and S6E and F). Remarkably, this revealed a subpopulation of high TREM2^+^/GFP^+^ CPC cells in Regimen-E relapsed tumours. Concurrent immunofluorescence and FISH analysis confirmed that these high TREM2^+^/GFP^+^ cells were indeed CPC cells with aberrant gains of chromosome 15 (Figure 5H). A similar but smaller and non-significant peak was also observed in radiation relapsed tumours. Thus, RECODR identified remarkable plasticity in malignant choroid lineages that underpins various modes of treatment relapse. In the context of combination AZD1390 and radiation treatment, this includes the outgrowth of progenitor-like CPC cells and a distinct, immune-like subpopulation of CPC cells.

We also detected an increase in (GFP^-^/TREM2^+^/CD45^+^/CD68/P2Y12^+^/CD49d^+^) tumour associated macrophages in Regimen-E relapsed CPC that could be readily distinguished from TREM2^+^ tumour cells (Figures 5G and I). Since tumour associated macrophages have been implicated in treatment resistance, these cells may have also contributed to Regimen-E resistance. Compatible with the observation that macrophages are the dominant immune cell type in the normal CP, we observed very few microglia in relapsed and control treated CPC^32^.

### RECODR predicts targets of dasatinib as a combination therapy resistance target

Similar to our analysis of monotherapy resistant tumours, we used RECODR ‘target scores’ to rank all genes in the R^RegE^-GN as potential mediators of treatment resistance (Figure 6A; Table S21). *Parp1* that appeared in the top 5% of AZD1390 and radiation monotherapy resistance targets fell to 405^th^ place in Regimen-E resistance targets, suggesting *Parp1* inhibition would not be a viable target for second line treatment in the context of combination therapy. However, RECODR ranked the transcripts of 12 targets of the multi-kinase inhibitor dasatinib above *Parp1,* with seven appearing in the top 2.5% (*Lyn, Syk, Hck, Vav1, Irak2, Fgr, Sla*; Figure 6A). The transcripts of the top five ranked targets were expressed to especially high levels in *Trem2^+^* immune-like CPC cells in Regimen-E relapsed tumours (Figure 6B). Furthermore, their immediate GN neighbours included 31% (1,808/5,849) of all genes on the R^RegE^-GN (Figure 6C). GFP/HCK (but not GFP/LYN) co-immunofluorescence analysis detected a subpopulation of high expressing immune-like CPC cells very similar to that by RECODR immune community enrichment scores and TREM2 expression in Regimen-E relapsed tumours (Figure 6D to G). In stark contrast, DEA identified only three of the 12 dasatinib targets in Regimen-E resistant immune-like cells relative to all other cells: these ranked outside the top 33% of differentially expressed genes (Figure 6H; Table S22). Thus, RECODR predicted that Regimen-E, but not AZD1390 or radiation treatment alone, drove the expansion of a subpopulation of immune-like cells that display myeloid mimicry that might be vulnerable to dasatinib therapy.

**Figure 6.**
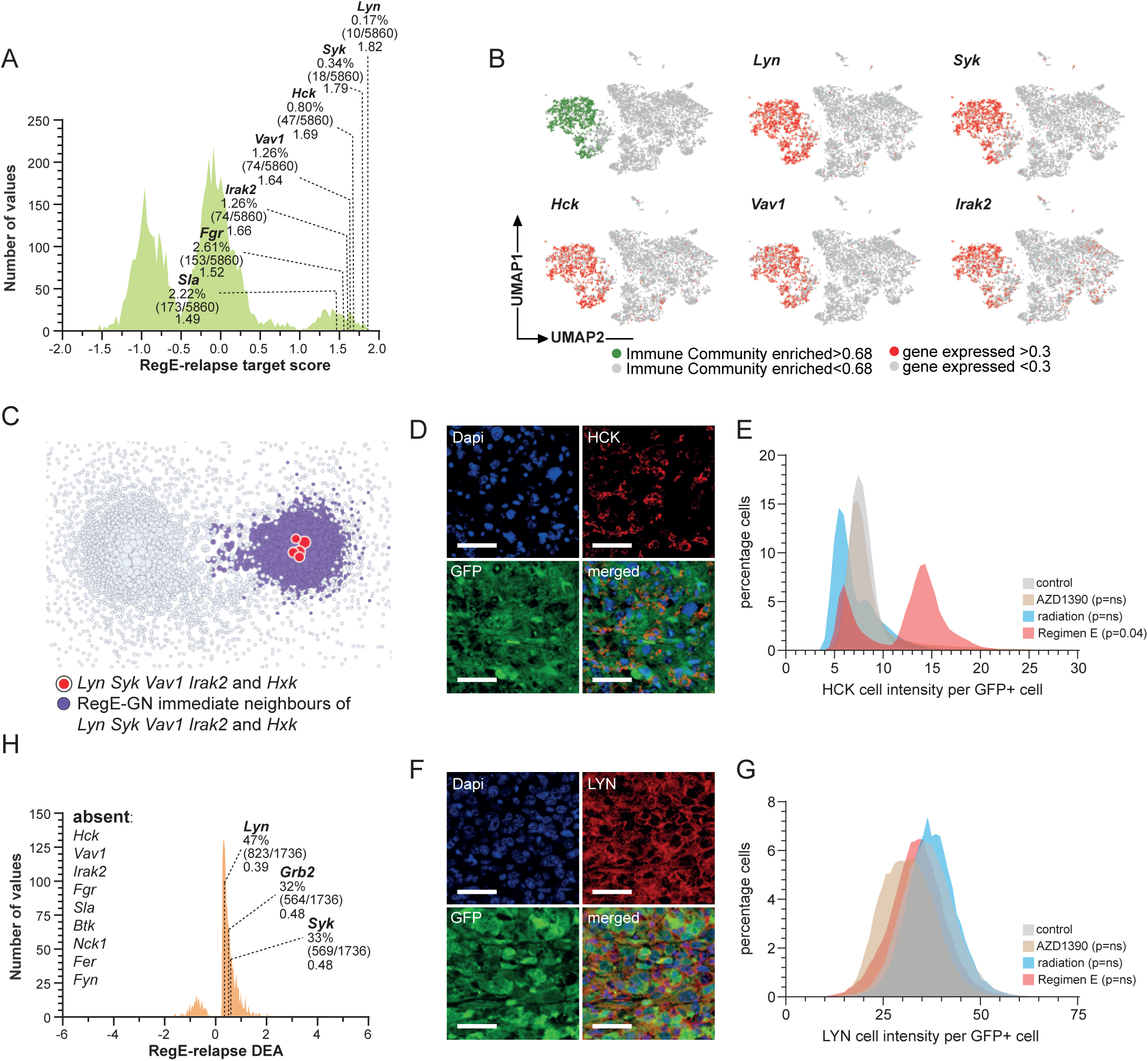
RECODR identification of drug targets in Regimen-E relapsed tumours. **A.** Frequency plot of ‘target scores’ of genes on R^RegE^-GN. The percentage ranking and target score is shown for dasatinib target genes. **B.** tSNE plots indicating immune community enrichment score (green top left) and dasatinib target gene expression (remaining plots) in scRNAseq profiles of Regimen-E relapsed CPC. **C.** R^RegE^-GN graph network showing *Lyn, Syk, Hck, Vav1* and *Irak2* (red) and their immediate neighbors (purple). **D.** Co-Immunofluorescence of HCK and GFP (Scale bar=40µm). **E.** Frequency plot of HCK immunofluorescence intensity/GFP^+^ CPC cells across different conditions and control (untreated CPC). **F.** Co-Immunofluorescence of LYN and GFP (Scale bar=40µm). **G.** Frequency plot of LYN immunofluorescence intensity/GFP^+^ CPC cell across different conditions and control (untreated CPC). **H.** Frequency plots of significantly differentially expressed (FDR<0.05) genes between immune-like cells in Regimen-E relapsed CPC versus immune-like cells in control CPC. Only *Lyn, Grb2* and *Syk* were significantly differentially expressed at a log-fold change of 0.25 or above: their percentage ranking is shown.

### Dasatinib prevents Regimen-E resistance

To test this if dasatinib might prevent CPC resistance to Regimen-E, we treated mice with CPC with relatively short (Regimen E-D^1^) or long (Regimen E-D^2^) durations of Regimen-E followed by continuous dasatinib treatment (25mg/kg/day; Figure 7A to C). We predicted that Regimen E-D^1^ would promote limited outgrowth of immune-like tumour cells and therefore would exhibit a decreased response to treatment with dasatinib, while Regimen E-D^2^ would promote more extensive outgrowth of immune-like tumour cells and therefore elicit a greater response to dasatinib. Remarkably, in keeping with this notion, Regimen E-D^1^ did not significantly improve either tumour growth suppression or survival relative to mice treated with Regimen-E or dasatinib alone (Figure 7A). In stark contrast, Regimen E-D^2^ significantly improved tumour growth suppression and mouse survival relative to control, dasatinib-only, or Regimen-E only therapy (Figure 7B). Notably, a modification of Regimen E-D^1^ in which AZD1390 and dasatinib therapy were alternated every two weeks also significantly suppressed tumour growth and extended mouse survival beyond that achieved by dasatinib-only or Regimen-E only therapy (Figure 7B).

**Figure 7.**
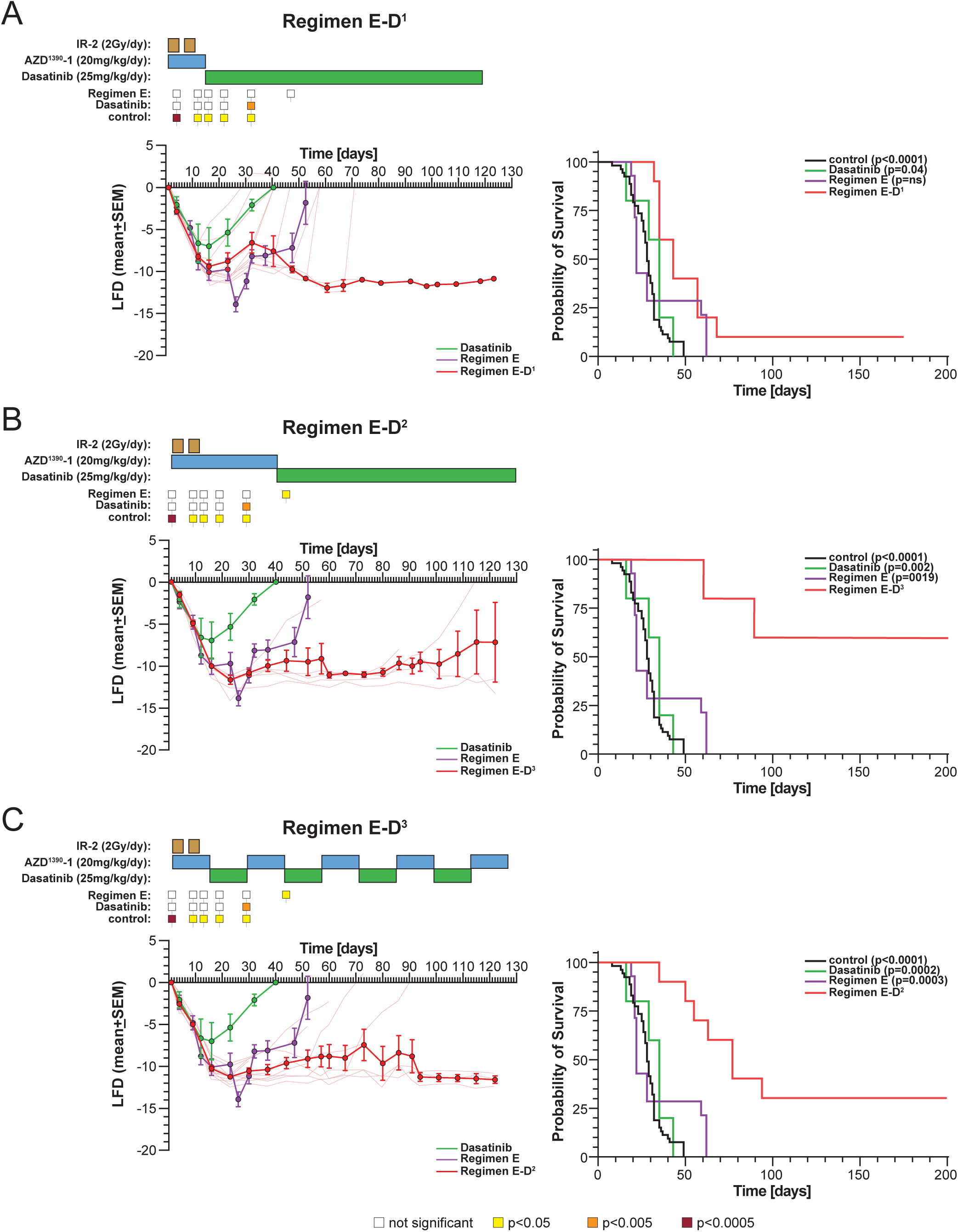
Dasatinib mitigates Regimen-E resistance. Graphs reporting (left in each) log fold difference growth (LFD) growth suppression (±standard error of mean) of treated relative to control treated mice with CPC and (right in each) survival of mice with CPC on the indicated treatment. Regimen E-D^1^ **(A)**, Regimen E-D^2^ **(B)** and Regimen E-D^3^ **(C)**. In growth suppression graphs coloured squares report significance (Wilcoxon adjusted p-value) of treatment-induced growth suppression relative to controls in plots (key at bottom of Figure). P values for each survival curve report the log-rank statistic relative to control treated mice.

## DISCUSSION

Drug resistance and treatment failure are responsible for the great majority of cancer deaths. Efforts to circumvent this resistance have focused on combining effective monotherapies in the hope that they will exhibit non-overlapping mechanisms of toxicity and resistance, thereby avoiding treatment failure. However, combination therapies also often fail through mechanisms that are hard to predict. Advanced computation is being used increasingly to improve the accuracy and efficiency of combination cancer therapy selection. This has included the use of network-based^52–54^, machine learning^55,56^, and deep learning approaches. While these approaches have the power to identify potential regulatory networks in disease, they typically select treatment targets by comparing patterns observed in the cells or tissues under study with curated pathways and interactions previously defined in other contexts. This approach has the caveat that selected targets may not perform the same function in the disease of interest, even if the associated gene modules are expressed.

Here, we introduce RECODR: a pipeline that combines co-expression-based GNs and NLP to track context shifting of genes within transcriptomes across cancer development and drug treatment conditions. This approach is based on the notion that co-expression patterns between genes across 1000s of cells can approximate the biological ground state of a tissue. To overcome the limitation of selecting potential drug targets based on their positioning within pathways or interaction states previously defined in other contexts, RECODR measures the magnitude of drift of the co-expression state of each gene between tissue states e.g., before and after treatment resistance. The operator can then use this as a measure of change of biological function. While the GNs generated by RECODR do not highlight direct interactions or causal relationships, they do scale, allowing the use of the entire transcriptome for each condition in single-cell datasets. This has the advantage of recording biological relevance within the context of the actual and entire transcriptome under study. This is likely to be especially relevant in cancer in which transient and fluid gene interactions can determine shifts in malignant phenotype^9,10^.

The detection by RECODR of a common mode of CPC resistance to AZD1390 or radiation monotherapy that differed when these treatments were combined, highlights the difficulty in predicting combination treatment resistance mechanisms. But it also illustrates the potential value of RECODR to unmask and mitigate combination therapy resistance mechanisms. Although we used RECODR to study an especially rare childhood brain tumour, the approach is disease type agnostic, giving it the potential to be utilized for other types of malignancies and disease pathologies.

Large-scale changes in gene context detected by RECODR during treatment resistance were driven by, and therefore had value in detecting underlying changes in, tumour cell subpopulations. In our study, RECODR revealed remarkable plasticity in CPC lineages. The GFP^+^/RFP^+^ cells that populate our CPC mouse model included progenitor, immune and mesenchymal-like cells that shared the same chromosomal CNVs; strongly supporting the notion that these cells share a common, clonal origin. Since lineage tracing of eCP has revealed a common origin for ciliated epithelial and neuronal cells^32^, then we predict this plasticity is inherited by daughter tumours, and that this plasticity generates immune-like CPC cells. Future work will be required to understand the functional implications of this plasticity, uncover additional treatment targets, and confirm our prediction that progenitor-like tumour cells sit at the apex of the CPC hierarchy.

Since CPC resisted both AZD1390 and radiation through the outgrowth of progenitor-like tumour cells, then the outgrowth of TREM2^+^ myeloid-like tumour cells following combination treatment resistance was unexpected. Immunosuppressive myeloid cells are established contributors to tumour progression and potential cancer treatment targets^49,57^. While most attention has focused on such tumour infiltrating immune cells, cancer cells can adopt myeloid features to create an immunosuppressing tumour microenvironment^48^. In CPC, the vast majority of TREM2^+^ cells that outgrew Regimen-E treatment were myeloid-like tumour cells. Notably, SYK–a dasatinib target that RECODR predicted could be targeted to mitigate combination treatment resistance–is a central to TREM2 signaling^51^. Furthermore, TREM2 is known to signal via SYK to promote myeloid cell survival in the brain and can be targeted to remodel the landscape of tumor-infiltrating macrophages^58^. Thus, dasatinib may target a TREM2-SYK signaling axis in myeloid-like CPCs to prevent their outgrowth and Regimen-E treatment failure. Review of combination AZD1390 and radiation clinical trials in glioblastoma and CPC will be important to determine if similar treatment resistance and mitigation mechanisms operate in these diseases.

## ACKNOWLEDGEMENTS

This work was supported by grants to Ri.J.G: Cancer Research UK (CRUK) Centre, CRUK Children’s Brain Tumour Centre of Excellence, CRUK Cambridge Institute Core Award; CRUK RadNet; the Brain Tumour Charity Quest for Cures. GB lab is supported by the UK Dementia Research Institute [award number UKDRI-2006] through UK DRI Ltd, principally funded by the UK Medical Research Council. N.D is supported by a Glenn/AFAR postdoctoral fellowship, Reagan Sloane Shanley research internship, William Randolph Hearst fellowship, and OFD/BTREC/CTREC Faculty Career Development Fellowship.

## AUTHOR CONTRIBUTIONS

BVN and ST conducted the bulk of the experimental procedures. EN, KEM, LR, MD, EC, LPH, GK, MA, Re.J.G, CT, GB, SPJ conducted/advised on important experimental procedures. MGF, AH, AP, ND, AR, and MKL provided important data and reagents. Ri.JG conceived the research and with AJ designed the approach and oversaw the research. All authors contributed to the writing of the manuscript.

## SUPPLEMENTARY FIGURE LEGENDS

**Figure S1.**
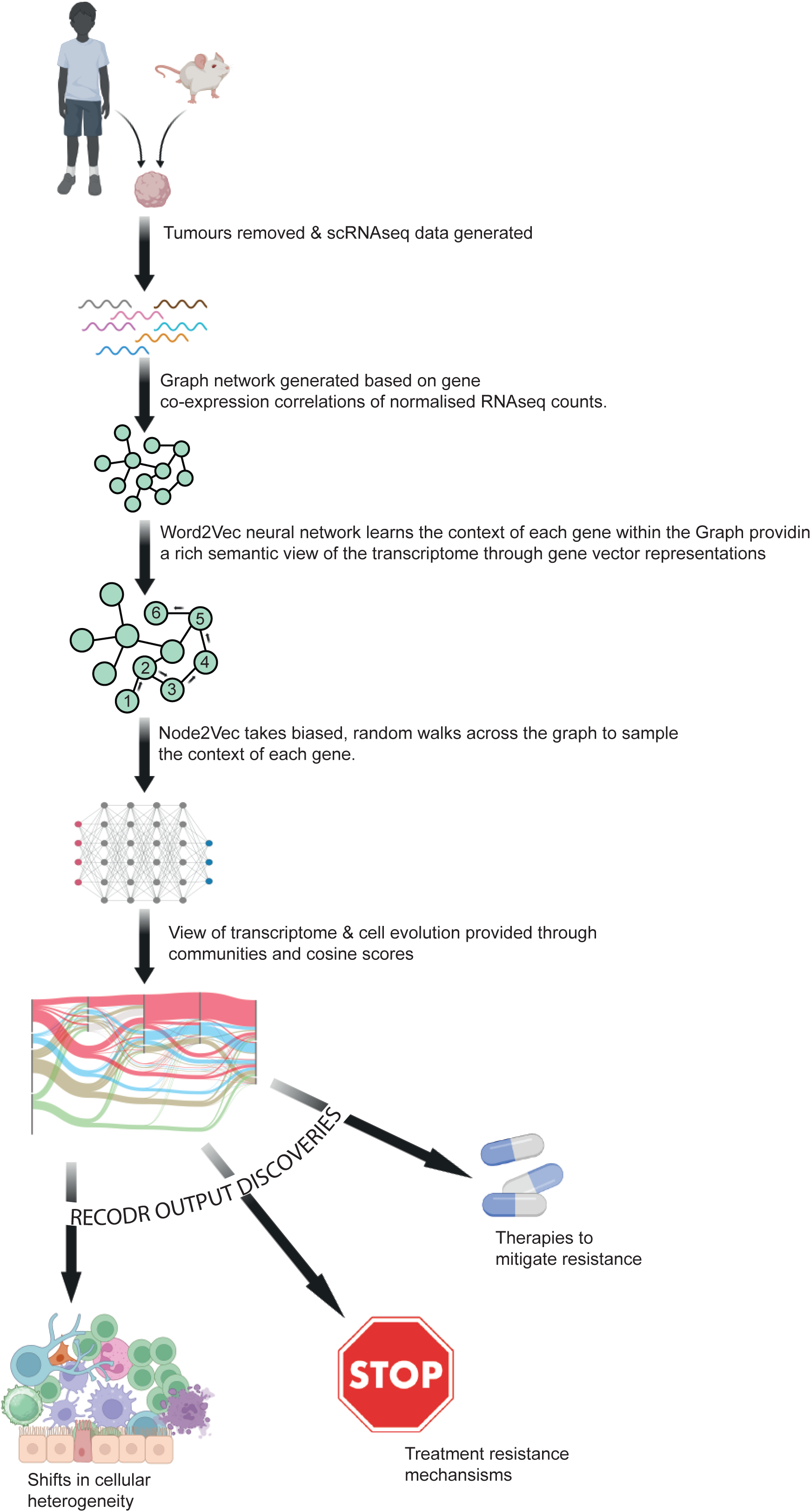
Summary of the RECODR pipeline.

**Figure S2.**
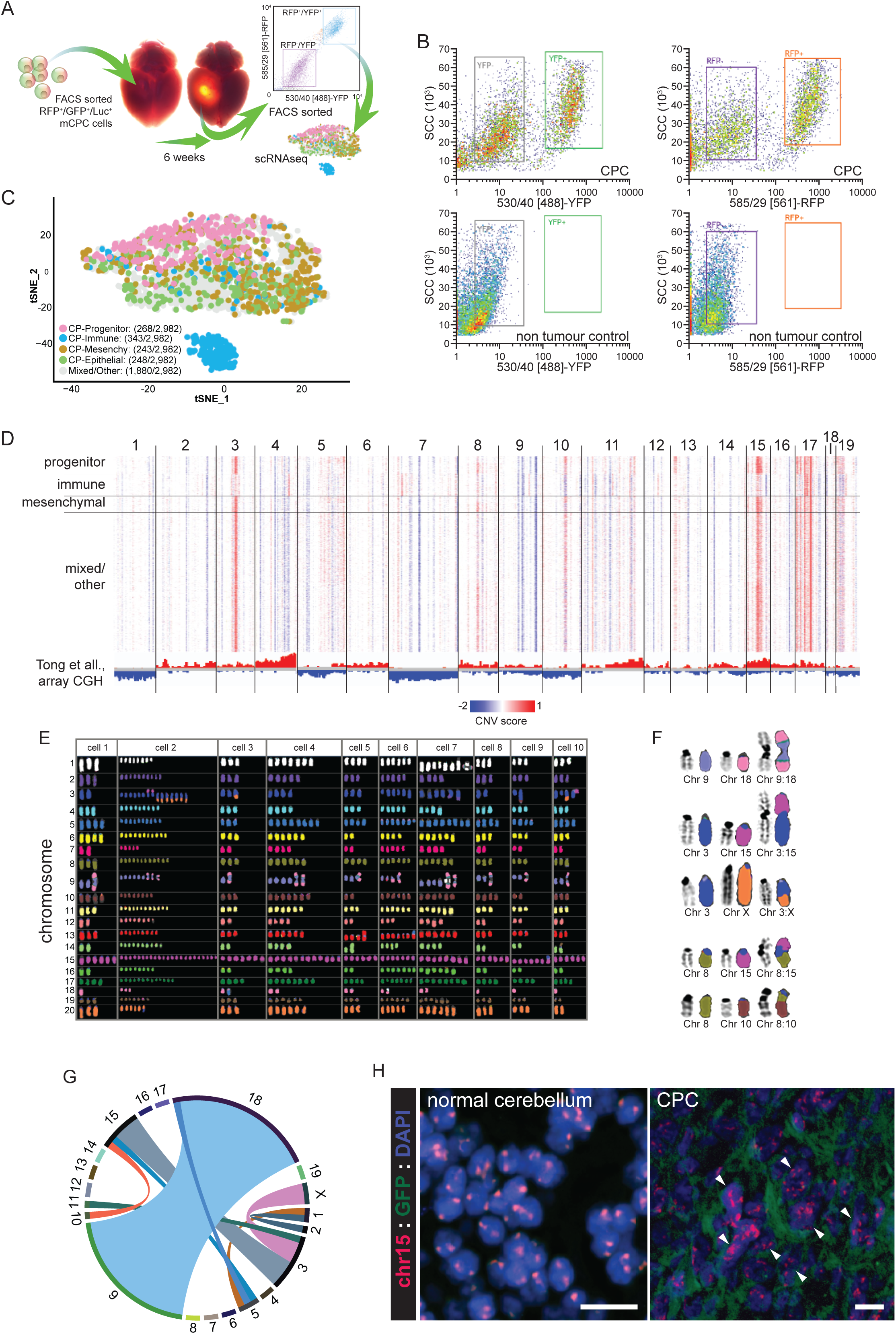
Isolation and transcriptome and DNA copy number profiling of mouse CPC. **A.** Schematic depicting CPC implants and FACS sorting for single-cell RNA sequencing (scRNAseq). **B.** Flow cytometry plots indicating gating of cells of YFP^+^ (left) and RFP^+^ (right) cells from tumour bearing (top) and non-tumour bearing (bottom) controls. **C.** tSNE of scRNAseq profiles of untreated CPC cells. Cells labelled according to enrichment of the indicated previously reported eCP cell type (Table S1; reference 32). **D.** Copy number variations (CNVs) inferred in scRNAseq profiles of each CPC cell in (C). CNV scores at bottom are from reference 30. **E.** Spectral karyotype analysis of 10 individual CPC cells. **F.** Representative SKY images of intracellular translocations in CPC cells. **G.** CIRCOS plot summarizing intracellular chromosomal translocations detected by SKY in 30 CPC cells. **H.** Concurrent GFP immunofluorescence and chromosome 15 FISH of the normal cerebellum (left) and CPC (right). Scale bar=20μm.

**Figure S3.**
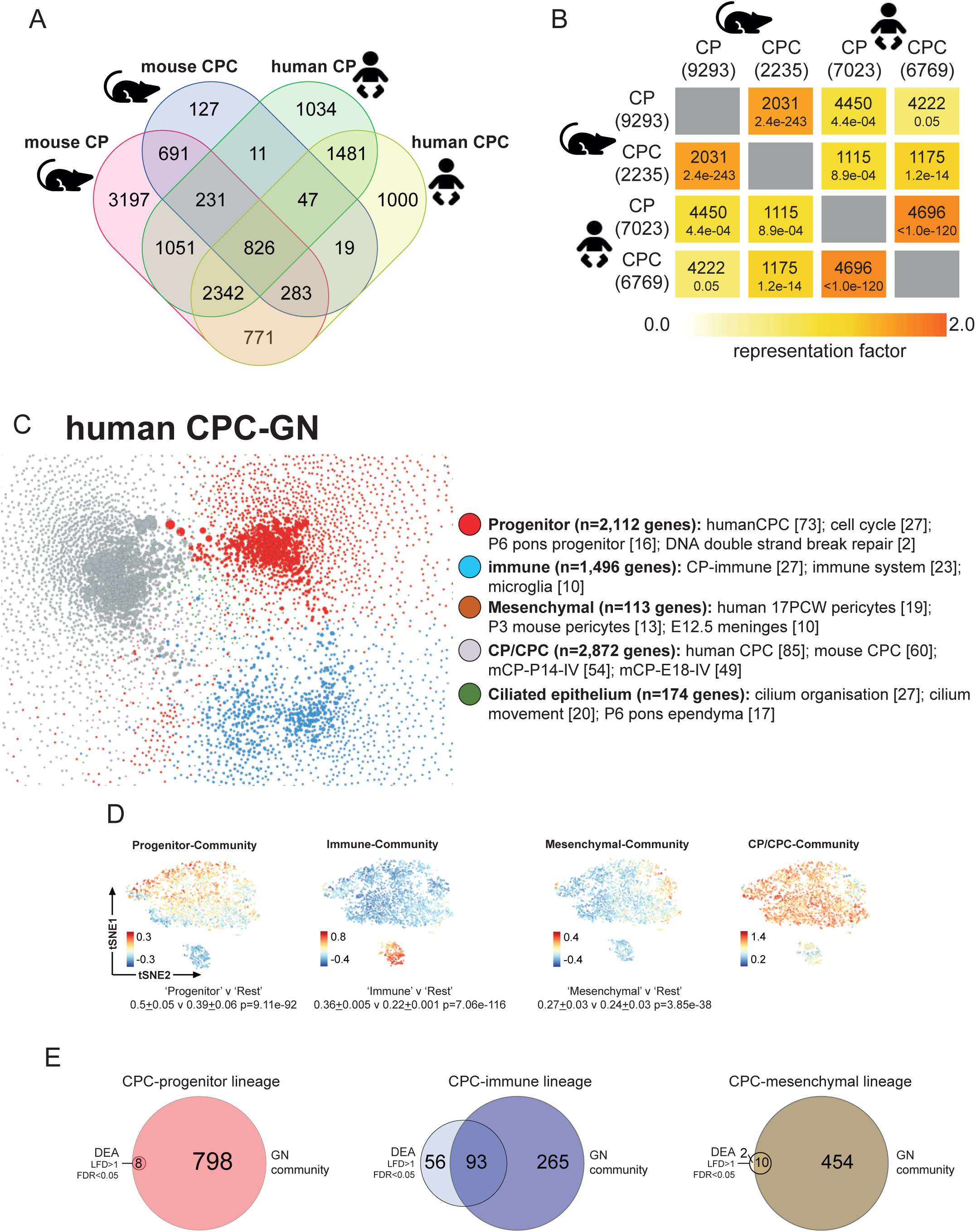
RECODR generated graph networks of mouse and human CPC. **A.** Venn diagram of overlap in genes among the indicated graph networks. **B.** Similarity matrix reporting the representation factor (top in each) and associated p-value (below in each) of overlap of genes between the indicated mouse and human GNs. **C.** R^CPC^-GN of human CPC with communities and associated enriched gene sets (Table S5). **D.** t-distributed Stochastic Neighbor Embedding (tSNE) plots of the indicated community gene signature enrichment in CPC scRNAseq profiles. **E.** Venn diagrams of overlap in genes detected in by GNs or differential expression in the indicated community and corresponding previously defined cell-type (according to reference 32).

**Figure S4.**
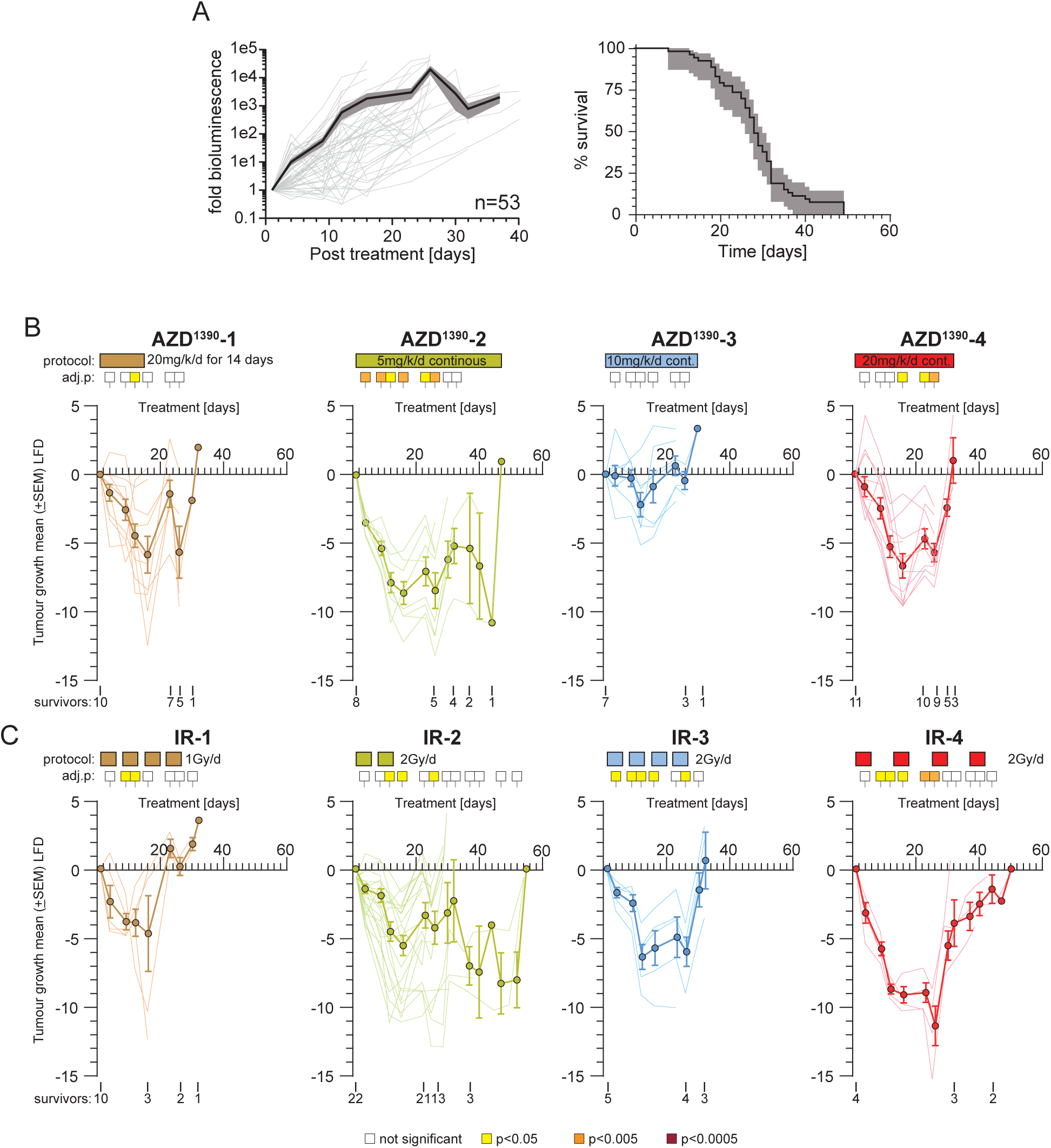
Mouse survival and/or relative tumour growth in mice with CPC. **A.** Characterisation of control treated mouse tumour growth (measured by bioluminescence, left) and mouse survival (right) in 53 mice with CPC. Different regimens of monotherapy AZD1390 **(B)** or radiation **(C)**. Coloured squares report significance (Wilcoxon adjusted p-value) of treatment-induced growth suppression relative to controls in plots below. (Bottom) Graphs of the average log fold difference (±standard error of mean) of each treatment relative to control treated CPC. Number of surviving mice at the indicated times each treatment are shown below each graph.

**Figure S5.**
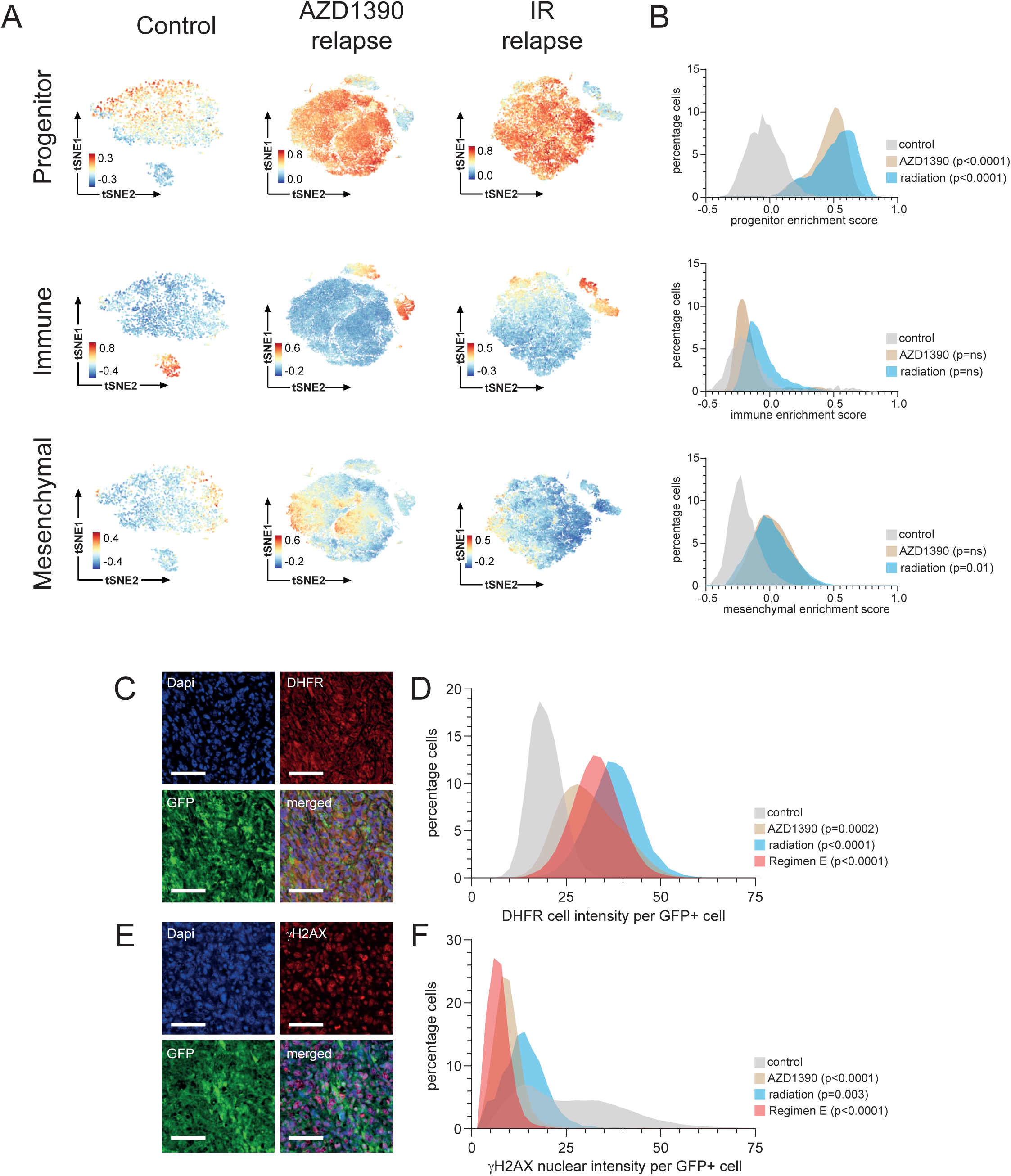
RECODR predicted changes in cell populations during monotherapy resistance. **A.** t-distributed Stochastic Neighbor Embedding (tSNE) plots of the indicated community gene signature enrichment score in scRNAseq profiles of cells isolated from the CPC treated as shown. **B.** Frequency plots of the progenitor (top), immune (middle) and mesenchymal (bottom) community enrichment scores in scRNA seq profiles across the indicated treatments. **C.** Co-Immunofluorescence of DHFR and GFP in exemplar radiation resistant CPC (Scale bar=40µm). **D.** Frequency plot of DHFR immunofluorescence intensity/GFP^+^ CPC cell across different conditions and control (untreated CPC). **E.** Co-Immunofluorescence of γH2AX and GFP in exemplar AZD1390 resistant CPC (Scale bar=40µm). **F.** Frequency plot of γH2AX immunofluorescence intensity/GFP^+^ CPC cell across different conditions and control (untreated CPC).

**Figure S6.**
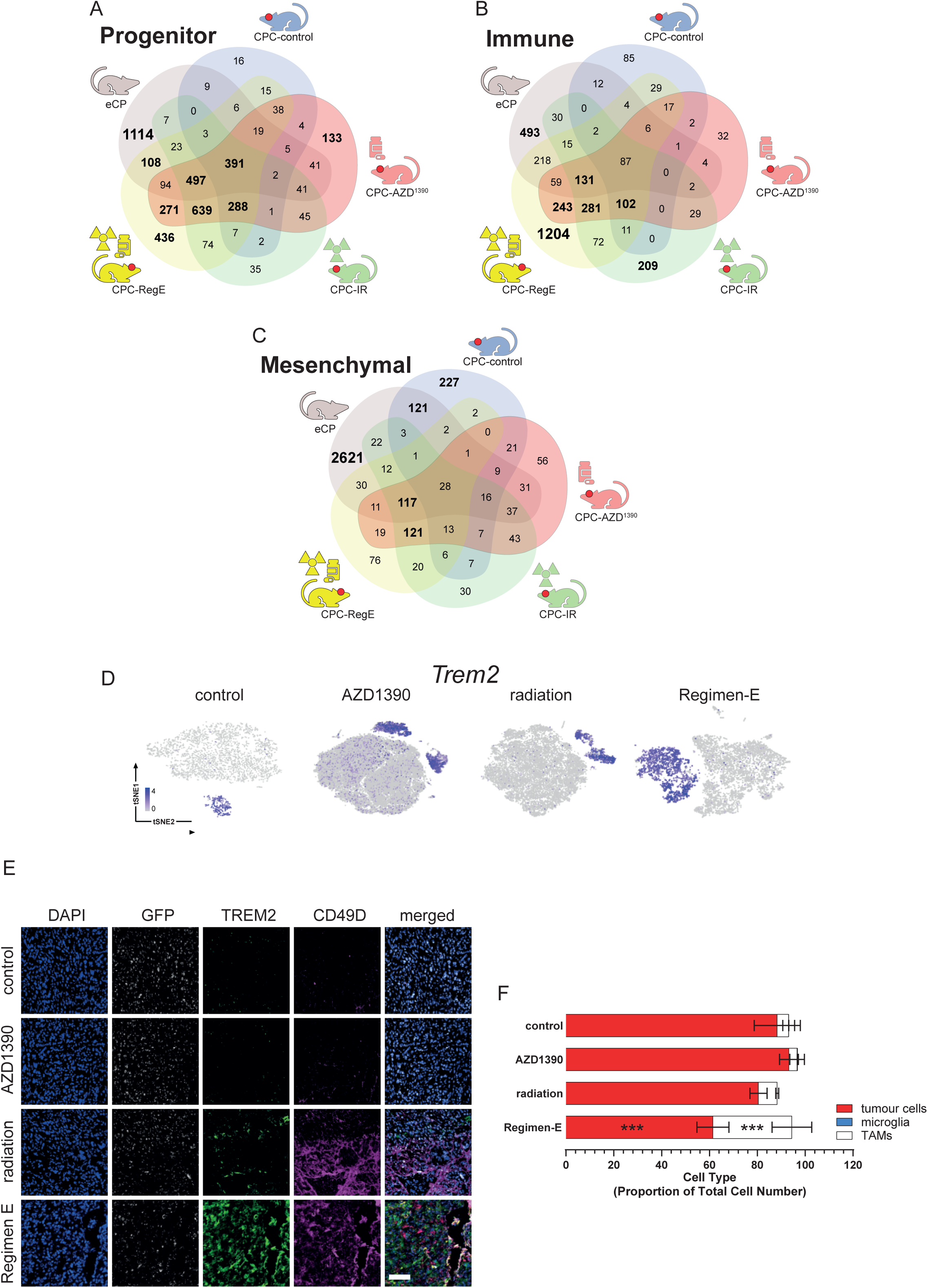
RECODR identification of transcriptome and cell population evolution following Regimen E resistance. Venn diagrams of gene overlap among progenitor **(A)**, immune **(B)** and mesenchymal **(C)** communities in the indicated conditions. **D.** tSNE plots of Trem2 expression in scRNAseq profiles of the indicated tumours. **E.** Co-immunofluorescence of GFP, TREM2, and CD49D in CPCs treated as shown. Scale bar=50μm. **F.** Mean proportion (±standard deviation) of tumour associated macrophages (TAMs; CD45^+^/CD68^+^/P2Y12^+^/CD49+), microglia (CD45^+^/CD68^+^/P2Y12^+^/CD49^-^) and GFP^+^ tumour cells across all conditions. ***, p<0.0005 Mann Whitney.

## STAR methods

### Cell Culture

The mc300 cell line derived from was maintained in cell culture at 37°C with 5% CO_2_. mc300 cells were maintained in neurobasal media (ThermoFisher scientific, 21103-049) supplemented with N2 (Gibco, 17502-048), serum-free B27 (Gibco, 17504044), L-Glutamine (Gibco, 25030-081) and penicillin/streptomycin (Gibco, 15140-122), 100µg/ml bFGF (Miltenyi Biotech, 130-093-243), 100µg/ml hrEGF (Miltenyi Biotech, 130-097-751) and 7.5% BSA (Sigma-Aldrich, A8412). Cells were grown until neurospheres formed. For cell passaging, neurospheres were centrifuged at 50g before being passed through a 40µm filter. Flasks were then rinsed with 5ml PBS and passed through the same filter. 2ml of StemPro Accutase (Gibco, A1110501) was then added and cells dissociated at 37°C for 15 minutes. Neurospheres were then triturated and passed through a 40µm filter into 20ml of media. Cells were then centrifuged at 118g for 5 minutes before being re-suspended in complete media at a 1:2 dilution.

### shRNA screen

shRNAs were generated and used in knock-down studies as described previously^59^. Briefly, shRNAs targeting each candidate DNA repair enzyme (or scrambled) were designed using the shRNA sequence prediction algorithm from Dharmacon/Thermo Scientific. shRNAs were cloned into the pFUGWH1-CFP vector and transformed into bacteria as one ligation product. The transformed bacteria were then spread on an LB/AMP plate and individual colonies were screened for unique shRNA constructs by sequencing. All constructs were sequence verified. DNA was fugene (Roche) transfected into 293FT cells (Invitrogen) along with lentiviral packaging plasmids for viral production. Virus titre was determined by cyan fluorescence protein (CFP) expression and transduced into CPC, RTBDN-ependymoma or mouse neural stem cells. Cells were then plated as single cells and colony formation determined after 72-120 hours.

### Chromosome painting

mc300 cells were plated in 6-well plates (1.5million cells/well) and incubated for 24 hours. Colcemid (Invitrogen, 15212012) was added to arrest cells in metaphase and incubated for 6 hours. After 6 hours, cells were triturated to dissociate clumps, collected in 1ml PBS and centrifuged at 400g for 8 minutes at room temperature. 10ml of hypotonic solution (0.0075M KCl in dH2O) was added, and cells were incubated for 12 minutes. After 12 minutes, 1ml of fixative (3:1 methanol: acetic acid) solution was added, and cells were centrifuged at 400g for 8 minutes at room temperature. Following centrifugation, the supernatant was removed, and 5ml of fixative solution was added and centrifuged at 400g for 8 minutes at room temperature. This washing step was repeated three times followed by dropping onto slides. The slides were allowed to dry for 3 hours. Metaphase was then confirmed by staining with DAPI (Sigma-Aldrich, D9542) followed by two washes with PBS and imaging by confocal microscopy.

For multiplex fluorescence in situ, hybridisation (M-FISH), 10µl of 24 colours human M-FISH paint probe mix generated at the Wellcome Trust Sanger Institute as previously described^60^ was denatured at 65°C for 10 min and then applied to the denatured slides. Slide denaturation was performed by immersing slides into alkaline denaturation solution (0.5M NaOH and 1M NaCl) for 40 seconds before being rinsed with 1M Tris-HCl at pH 7.4 for 3 minutes followed by 1X PBS for 3 minutes and then dehydrated with 70%, 90% and then 100% ethanol. For hybridisation, slides were incubated for 10 minutes at 37°C for 40-44 hours. After incubation, slides were washed with 0.5X SSC for 5 minutes at 75°C, then rinsed with 2X SSC+0.05% Tween 20 for 5 minutes and washed with 1X PBS for 2 minutes at room temperature. Slides were then mounted with SlowFade® Diamond Antifade Mountant containing DAPI. Images were taken on a Zeiss AxioImager D1 fluorescent microscope. M-FISH images were captured with the SmartCapture software (Digital Scientific UK) and processed with SmartType Karyotypes software (Digital Scientific UK).

### Animal work

All animal work was carried out under the Animals (Scientific Procedures) Act 1986 in accordance with the UK Home office license (Project License PP9742216) and approved by the Cancer Research UK Cambridge Institute Animal Welfare and Ethical Review Board. Mice were housed in individually ventilated cages with wood chip bedding plus cardboard fun tunnels and chew blocks under a 12 hour light/dark cycle at 21± 2°C and 55%± 10% humidity. Standard diet was provided with ad libitum water. All mice were housed for habituation for at least 1 week before the start of the experiment. The Crl:CD1-*Foxn1nu, 086* mouse strain was used. Mice for all experiments were between 7-9 weeks old at the start of the experiment. Mice were orthotopically implanted with 5000 cells of the mc300 cell line in 5μl of matrigel (Corning, 354230) per mouse. mc300 cells below passage 14 were used for tumor implants. Tumor growth was monitored by bioluminescent imaging (BLI). 3 days post implant, the tumors reached a suitable signal of 5*10^5^ p/s/cm2/sr (BLI) and animals were then randomized into control and experimental groups. Mice were continually monitored for clinical signs and tumor progression was measured by BLI. For BLI monitoring, mice were intraperitoneally (IP) injected with 15mg/ml of D-Luciferin (Perkin-Elmer, 122799) and placed into a housing chamber containing isoflurane. Once anesthetized, mice were placed into the IVIS spectrum and positioned into a nose-cone with continued delivery of isoflurane. BLI was then measured.

### Drug Preparation

AZD1390 (AstraZeneca) was diluted into a vehicle of water + 0.1% Tween 80 (Sigma-Aldrich, P1754) at 2mg/ml stock concentration. The drug was dissolved with a magnetic stirrer at room temperature and left to continually stir until administration. Dasatinib (MedChemExpress, HY-10181) was prepared in 80mM of citric acid monohydrate at 2.5mg/ml and manually mixed, then stored at 4°C. AZD9574 was prepared in deionized water in methanesulfonic acid (Sigma-Aldrich, 471356) at a pH of 3.0-3.2. 0.3mg/ml of AZD9574 (AstraZeneca) was diluted in the vehicle and manually mixed and stored at 4°C. All drugs were allowed to come to room temperature before administration.

### Treatment

#### Irradiation

Mice receiving radiation were anesthetized by isoflurane in a housing chamber before being put into the Small Animal Radiotherapy Research Platform (SARRP). Mice were positioned into nose-cones in the SARRP with continued isoflurane delivery. 20Gy of targeted radiation to the implant site was given to animals with 2Gy/day in cycles of 5 days on and 2 days off.

#### Drug treatment

For mice undergoing drug treatments, drugs were delivered by oral-gavage (Instech Laboratories plastic feeding tubes, 20ga x 38mm, Linton Instrumentation, FTP-20-38) daily according to treatment schedules outlined in this paper. AZD1390 was administered at a concentration of 20mg/kg, Dasatinib was administered at a concentration of 25mg/kg and AZD9574 was administered at a concentration of 3mg/kg.

### Tissue collection and processing for IHC

Once mice reached a humane clinical endpoint, mice were sacrificed by a rising concentration of CO_2_. Brains were harvested by immediate decapitation, posterior to the occipital bone followed by removal of the brain. Brains were fixed in 10% neutral buffered formalin for 24 hours, followed by 70% EtOH for a further 72 hours before being paraffin-embedded. For immunohistochemical studies, 7-μm thick sagittal sections were used.

### Immunofluorescence staining

Briefly, sections were first deparaffinized and treated using a standard antigen unmasking step in 10 mM Tris/EDTA buffer pH 9.0. Sections were then blocked with Mouse BD Fc Blocking solution (BD Biosciences, 553141) and treated with True Black Reagent (Biotium, 23007) to quench intrinsic tissue autofluorescence.

Detection of mouse chromosome 15 was performed on FFPE sections. Pretreatments were carried out using the Tissue Pretreatment Kit (OGT, LPS100) according to manufacturer’s instructions: Incubation with Tissue Pretreatment Solution (Reagent 1 at 98°C for 10 minutes followed by 3 minutes wash in milliQ water twice, 8 minute incubation with Enzyme Reagent (Reagent 2) at 37°C followed by 3 minutes wash in milliQ water twice). Slides were dehydrated through graded ethanol. Ready-to-use XMP 15 Mouse Chromosome painting probe (Metasystems, D-1415-050-OR) was applied to each slide and coverslips were sealed with Fixogum rubber cement (VWR, ICNA11FIXO0125). Slides were denatured for 5 min at 75°C before hybridisation overnight in a humid chamber at 37°C. Rubber cement was carefully peeled off and coverslips were removed by soaking in 2X SSC + 0.05% Tween 20 (Sigma-Aldrich, P1379) followed by a wash in 0.4X SSC buffer at 72°C for 2 min and 2X SSC + 0.05% Tween 20 for 30 seconds. Slides were incubated with DAPI for 5 mins at room temperature in the dark. Slides were washed for 5 minutes each in 3 changes of PBS prior to mounting with Prolong Diamond Antifade Mountant (ThermoFisher Scientific, 36961).

The sections were then treated with a 1-minute additional heat mediated antigen retrieval step in Tris/EDTA buffer. The sections were then immunoreacted for 1 h at RT using 1 μg/ml cocktail mixture of immunocompatible antibody panels (see Table 1 for antibody sources and technical specifications). This step was followed by washing off excess primary antibodies in PBS supplemented with 1 mg/ml fish skin gelatin (Sigma-Aldrich, G7041) and staining the sections using a 1 µg/ml cocktail mixture of the appropriately cross-adsorbed secondary antibodies (purchased from either ThermoFisher Scientific, Jackson ImmunoResearch or Li-Cor Biosciences) conjugated to one of the following spectrally compatible fluorophores: DyLight 405, Alexa Fluor 488, Alexa Fluor 546, Alexa Fluor 594, Alexa Fluor 647, Cy5.5 or IRDye 800CW. After washing off excess secondary antibodies, sections were counterstained using DAPI to visualize cell nuclei. Coverslips were then placed on slides using Prolong Diamond Antifade Mountant and imaged using the Vectra Polaris (described in ‘Image Acquisition’ below).

**Table 1.**
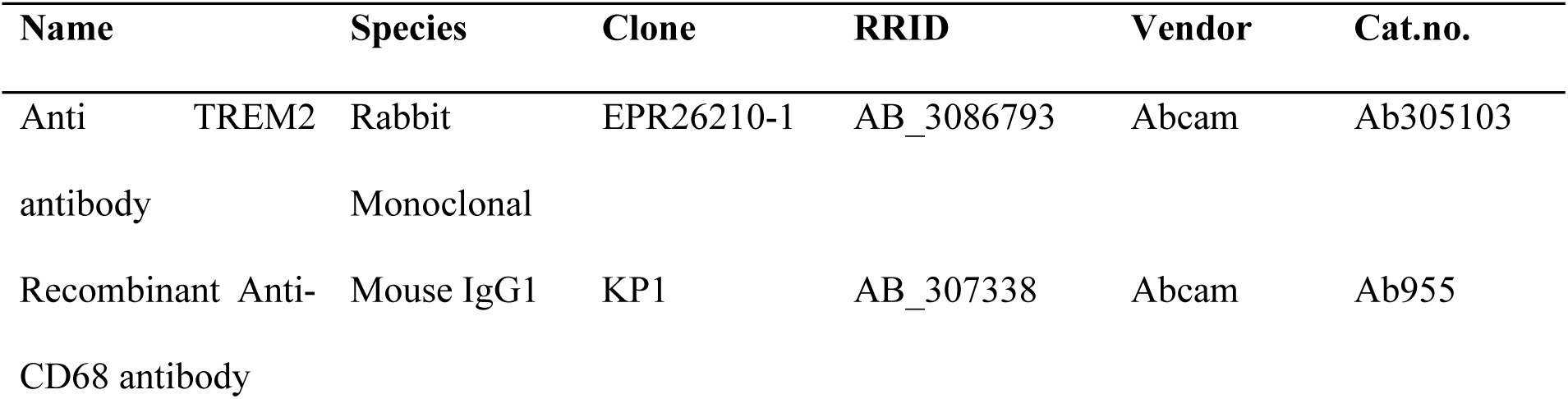
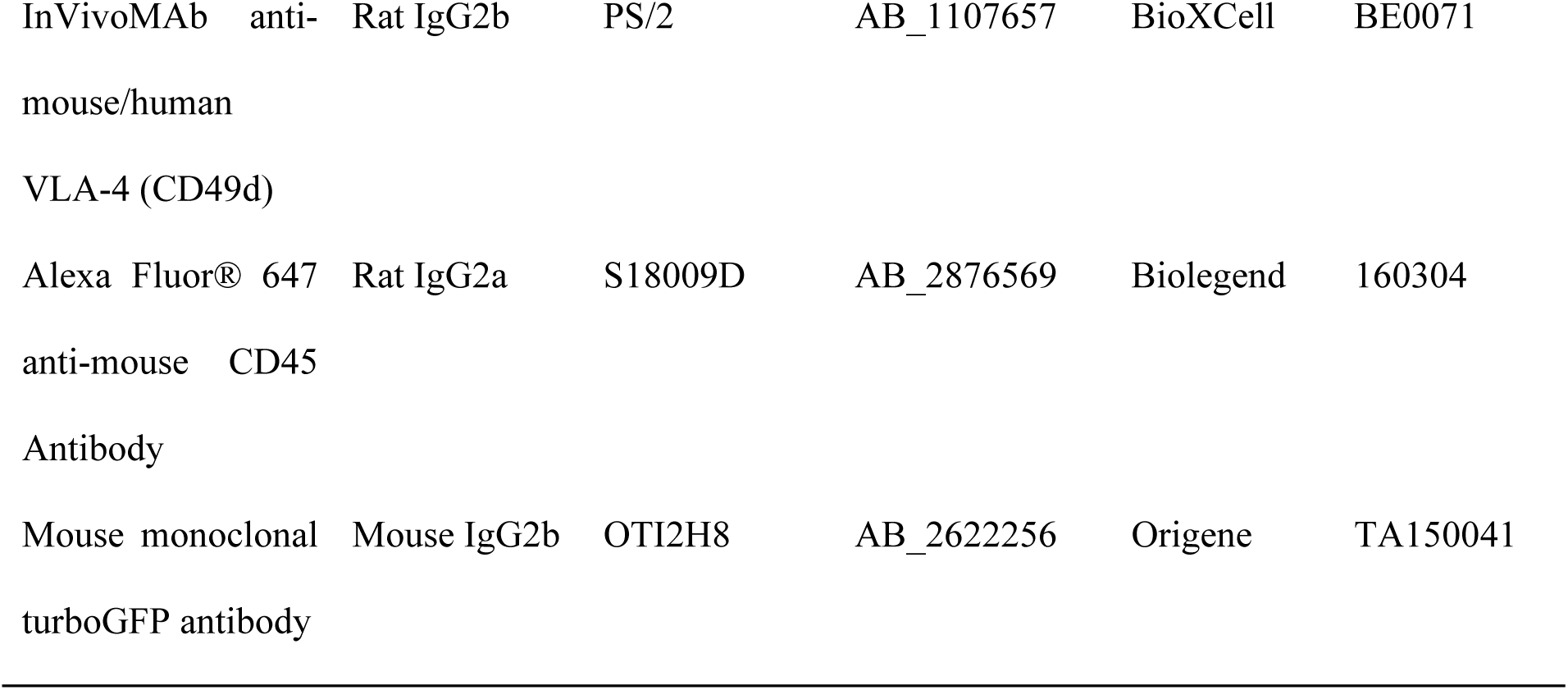
Primary antibodies used.

### Image Acquisition

All fluorescently labeled slides were scanned on the Vectra Polaris at 40X magnification using appropriate exposure times. Whole-slide images were scanned with 7-color whole-slide unmixing filters (DAPI + Opal 570/690, Opal 480/620/780 and Opal 520). Library slides were generated from representative tissue sections to allow for accurate unmixing of the multiplexed samples, including a slide stained for each single fluorophore, a DAPI only slide and an autofluorescence slide wherein no antibody, or DAPI was applied. The unmixing performance of this tissue-specific spectral library was compared to that of the synthetic Opal library available in inform. Resultant image tiles were then stitched together within HALO(R) (Indica Labs) to produce a whole-slide multichannel, pyramidal OME-TIFF image for downstream imaging analysis.

### Fluorescence Activated Cell Sorting (FACS)

Tumors were freshly isolated from mice by micro-dissecting and dissociation for one hour at 37°C in enzymatic dissociation solution containing 20U/ml of papain (Sigma-Aldrich, P4762) and 100µg/ml of DNAseI (VWR, A3778) in high glucose DMEM (Gibco, 11965092) with 2mM L-Glutamine, 5% penicillin/streptomycin, and 10% FBS (Gibco, A5256801). Following dissociation at 37°C, cells were triturated 5-6 times before being passed through a 40μm filter. The original tube containing cells was washed with 5ml of HBSS (ThermoFisher Scientific, 14175095) which was then passed through the same 40μm filter. Cells were then centrifuged for 5 mins at 4°C and 300g. Following centrifugation, the supernatant was discarded, and the cell pellet was re-suspended in 10ml HBSS before being passed through a second 40μm filter. This cell suspension was then centrifuged for 5 mins at 4°C and 300g. Following centrifugation, cells were re-suspended and transferred to a FACS tube. Dual RFP and YFP cells were flow sorted using a BD Aria II (BD Biosciences) with the following gating strategy, forward and side scatter, singlets and dual RFP (561nm-585/29) and YFP(488nm-530/40) positive cells. Dead cells were excluded with DAPI, UV (355nm-450/50).

### Single cell RNA-sequencing

Sorted cells were submitted for 10X sequencing and libraries were prepared using the Chromium Single Cell 3’ Reagents kit v.3. Briefly, samples were resuspended in PBS with 0.04% BSA and loaded onto the Chromium microfluidic chip(10X Genomics) to generate single-cell bead emulsions with individual 10X barcodes. RNA from barcoded cells was reverse transcribed in a C1000 Touch Thermal cycler (Bio-Rad) and libraries were prepared according to manufacturer’s protocols (12 cycles were used for cDNA amplification). cDNA quantity and quality was measured using an Agilent TapeStation 4200 (High Sensitivity 5000 ScreenTape). About 375ng of material was used for library preparation and library quality was assessed with a TapeStation 4200 (High Sensitivity D1000 ScreenTape). Libraries were normalized to equal molar concentrations (10nM) and pooled. Sample pools were sequenced on a NovaSeq 6000 to the following parameters: 28bp, read 1; 8bp, i7 index and 91bp, read 2, aiming for 20000 reads per cell. RNA reads were processed by Cellranger (10x genomics) and aligned to the mouse (mm10) genome. The filtered gene matrices were then used as the input for downstream analysis pipelines.

### Single cell/nucleus RNA-sequencing analysis

Single-cell RNA-sequencing was analyzed in R using Seurat. First, for each condition, individual samples were read into R using the *Read10X* function followed by merging samples from the same condition using the *merge* function. The *PercentageFeatureSet* function was then used to calculate the percentage of counts derived from mitochondrial genes. The data was filtered to include cells with more than 200 genes, less than 3000 genes and less than 20% mitochondrial content per cell. Next, cells were log normalized using the *NormalizeData* function. The *FindVariableFeatures* and *ScaleData* functions were then run to find the top 2000 most variable features and scale the dataset (with total number of RNA counts and percent mitochondria regressed out). Downstream clustering was then carried out by applying the *RunPCA*, *FindNeighbors*, *FindClusters*, *RunUMAP* and *RunTSNE* functions using default parameters. Differential expression between different cell types was carried out with the Seurat pipeline run on merged control and relapse setting datasets and then using the Wilcoxon Rank Sum test with the *FindMarkers* function with default parameters for Seurat version 4. Normal mouse choroid-plexus single-cell datasets were taken from GSE168704 and processed using the standard Seurat workflow. Human single nucleus RNA-sequencing (sNuc-seq) of human choroid and human CPC from DKFZ German Cancer Research Center were used. The appropriate ethics approval was acquired prior to sequencing and the data were processed using the standard Seurat pipeline. Enrichment of gene signatures in cells was carried out using the R package UCell^61^. Enrichment scores of gene signatures were added to the Seurat object using the *AddModuleScore_UCell* function.

### Inferred copy number variation analysis

To infer copy number variation (CNV), all Seurat objects generated in ‘Single cell RNA-sequencing analysis’ (including GSE168704) were merged using the *merge* function and the R library DropletUtils was used to generate the merged single cell dataset in the traditional cellranger format. The *write10xCounts* function of DropletUtils was used to generate this format using raw counts of the Seurat object. The remainder of the analysis was carried out in Python. The output of *write10xCounts* was read in as a single cell object in Scanpy using the *read_10x_mtx* function. Next, raw counts were log normalized using the *normalize_total* and *log1p* functions from Scanpy. Then, the infercnvpy package was used to infer CNV in our single-cell dataset. Following normalization, genomic positions were added to the Scanpy dataset using the *genomic_position_from_gtf* function with the gencode vM23 primary assembly. Next, the *infercnv* function was run with normal choroid plexus cells (3V, 4V and LV single-cells from GSE168704) as the reference category and default parameters. Clustering was run with infercnvpy with the *pca*, *neighbors* and *leiden* functions using default parameters. Plots were generated using the *chromosome_heatmap* function.

### RECODR

The RECODR pipeline was created to take expression data from two or more conditions and carry out co-expression networks of each condition with a natural language processing approach on top of the graphs to understand the context in which genes are found with respect to their co-expression patterns. An approximation of how the context of the same gene between different conditions changes is then carried out with model alignment and the cosine similarity measure. The constituents of the pipeline are described below.

#### Graph Networks and community detection

For each condition (including GSE168704 and sNuc-seq of human choroid and CPC), the log-normalized counts were taken from the Seurat objects from the ‘single cell RNA-sequencing section’ and converted to dataframes. Next, the dataframes were filtered to include only genes that were expressed in at least 5% of cells. A pairwise Pearson correlation analysis was carried out between every pair of genes using the Pandas *corr* function in Python to identify the extent of linear correlation between gene pairs. The correlation matrix was then stacked using the Pandas *stack* function to generate a source, target and correlation value edgelist. Gene pairs with a Pearson correlation coefficient of 0.1 or above were taken forward for subsequent analysis. Pearson correlations between the same gene were removed to prevent self-connections in the graph network. The NetworkX function *from_pandas_edgelist* was used to generate an undirected and unweighted graph network for each condition following the steps described above where nodes represent genes, and edges connect nodes if those genes have a Pearson correlation of 0.1 or above. Community detection was carried out by the *best_partition* function from the python-louvain library using default parameters to identify modules in our graph networks. For plotting, NetworkX graphs were saved to the Gephi format with the *write_gexf* function and then plotted using the freely available software Gephi.

#### Community enrichment

Analyzed Seurat objects representing each CPC condition were converted to ‘h5ad’ files and then read in as Scanpy objects in Python using the *read_h5ad* function. Community files for each condition were read in as CSV files using the Pandas *read_csv* function. Genes from each community were subset and the *score_genes* function from Scanpy was used to generate a per cell enrichment score of each community for cells in each condition.

### g:Profiler analysis of communities in graph networks

g:Profiler analysis of communities was carried out using the g:Profiler Python package incorporated into a newly built function that uses each list of genes in each community of a graph and runs the default g:Profiler pipeline, with or without a custom GMT file.

### Node2Vec and Word2Vec

To leverage the use of vector-based representations of nodes on each graph we employed Node2Vec followed by the skip-gram objective of the Word2Vec architecture to learn the graph structure for each condition.

#### Node2Vec

To represent the graph networks in a format that adequately represents node features, we used Node2Vec^38^. Node2Vec carries out biased random walks (second order random walks) that sample neighborhoods around nodes. Briefly, normalized transition probabilities were calculated for every node such that the likelihood of transitioning from one node to another is determined. The walks are influenced by two hyperparameters which determine the likelihood of returning to the node the walk originated from or going outward from the current node. These hyperparameters are the return parameter *p* which we set to 1 and the ‘in-out’ parameter q which we also set to 1. 200 walks were carried out from every single node on each graph for a walk length of 80 nodes. This random walk represents the graph context and generates the input format for Word2Vec.

#### Word2Vec

Word2Vec is a neural network which aims to understand the context in which words are found in text^39^, generating rich semantic representations of words. This is achieved through static vector representations of each word in a corpus of text where vectors are optimized such that words that are found in the same context have similar vectors and words that are not found in the same context have dissimilar vectors. The resulting word vectors describe the relationship of that node and all others around that node. The similarity between vectors can be measured by metrics such as the cosine distance. The Node2Vec library used internally calls on Word2Vec from the Gensim natural language processing library. We ran the skip-gram objective with negative sampling of the Word2Vec architecture with a vector *dimension* of 64, *window* size of 10, *min_counts* of 5, initial learning rate (*alpha)* of 0.025 and 5 negative words (*negative)*. All models were trained for 10 epochs.

### Alignment of Word2Vec models

Training of each Word2Vec model is inherently stochastic due to the random weight initialization step when models are created. For this reason, The same node on a graph cannot be compared to the same node on another graph immediately after training as the vector spaces will be different. To measure how each gene changes context as a function of treatment condition, we carried out the Procrustes alignment between the embedding matrices of our Word2Vec models in a pairwise manner. The alignment is an orthogonal transformation of an embedding matrix such that a word embedding matrix for one model is aligned to a target matrix from another model. This is carried out using only common words between both models. The aim is to find the orthogonal matrix that best maps to the target embedding matrix to which the model is being aligned allowing for comparisons of the same word across models. This is achieved by minimizing the sum of square distances between vectors and is solved by the singular-value decomposition^62^.

### Comparing vectors across models

Following Word2Vec model alignment, the cosine similarity was used as a measure of context drift for words between two models. To generate the cosine distance between two vectors, the *cosine* function in SciPy was used to compare the same gene in two different Word2Vec models. Cosine similarity was taken as one minus the cosine distance yielding a range of values between -1 and 1. For presentation, we calculated the vectors of the baseline against themselves, generating vectors equal to 1. For genes that are absent in either the baseline model or comparator model, we assigned a value of -1.1. Any genes present in the aligned model but not present in the baseline model were given a value of 1.2. Any value outside the range of -1 to 1 are indicator values and do not represent a metric as no vector arithmetic is carried out.

### Node target scores

To identify potential drug targets in the relapse setting we generated a node scoring system based on the context drift of genes from the NLP analysis plus characteristics of each gene on the graph network. For each gene on a graph in the relapse condition, a cumulative score was generated using five scoring criteria. The first criteria was the presence of a gene on a graph in the relapse setting versus the presence of that gene in the untreated tumor setting. If a gene was only found in the relapse graph, a score of 0.01 was given, and if the gene was also present in the untreated graph, a score of -1 was given. The second criteria was a neighbor score. For each gene, its edges were compared to the original CPC graph. The number of edges that were unique to the relapse setting were counted and normalized to the total number of nodes in the relapse graph. The third criteria was a graph reach score. As a proxy for how much of the graph a node could reach, we took the neighbors of a node plus the unique edges of the 2nd hop neighbors. The number of unique genes was normalized to the total number of genes in the relapse graph. The fourth criteria was context drift. The cosines with the relapse graph as the baseline were taken and compared to the untreated CPC Word2Vec model. Next, the cosine dataframe was subset to include only the neighbors of the node being analyzed. Genes that were not present in the untreated CPC graph were removed from the dataframe and the mean of the remaining cosines was taken as a measure of overall context drift. one minus the mean context drift was taken to reward higher context drift of genes in the relapse setting relative to the untreated CPC graph. Finally, a drug score was given where if a drug exists for a specific target according to the canSar.ai database, a score of 0.5 was given, and if a drug does not exist against a gene, a score of -1 was given. The sum of all scores was taken to generate a final score for each gene.

### Circos Plot

For chromosomal translocations, ‘recipient chromosome’ and ‘donor chromosome’ tables were created with the proportion of translocations observed. The *chordDiagramFromDataFrame* function from the Circlize package in R was then used to generate the circos plot.

### Alluvial Plots

Alluvial plots were generated using the open source software RAWGraphs 2.0.

### Value scaling

Plots indicating scaled-scores were scaled using Pythons sci-kit learn library with the *minmax_scale* function. Scaling was carried out across the ‘sample’ axis in order to ensure the feature range of values was scaled across the sample of interest.

### Software and algorithms

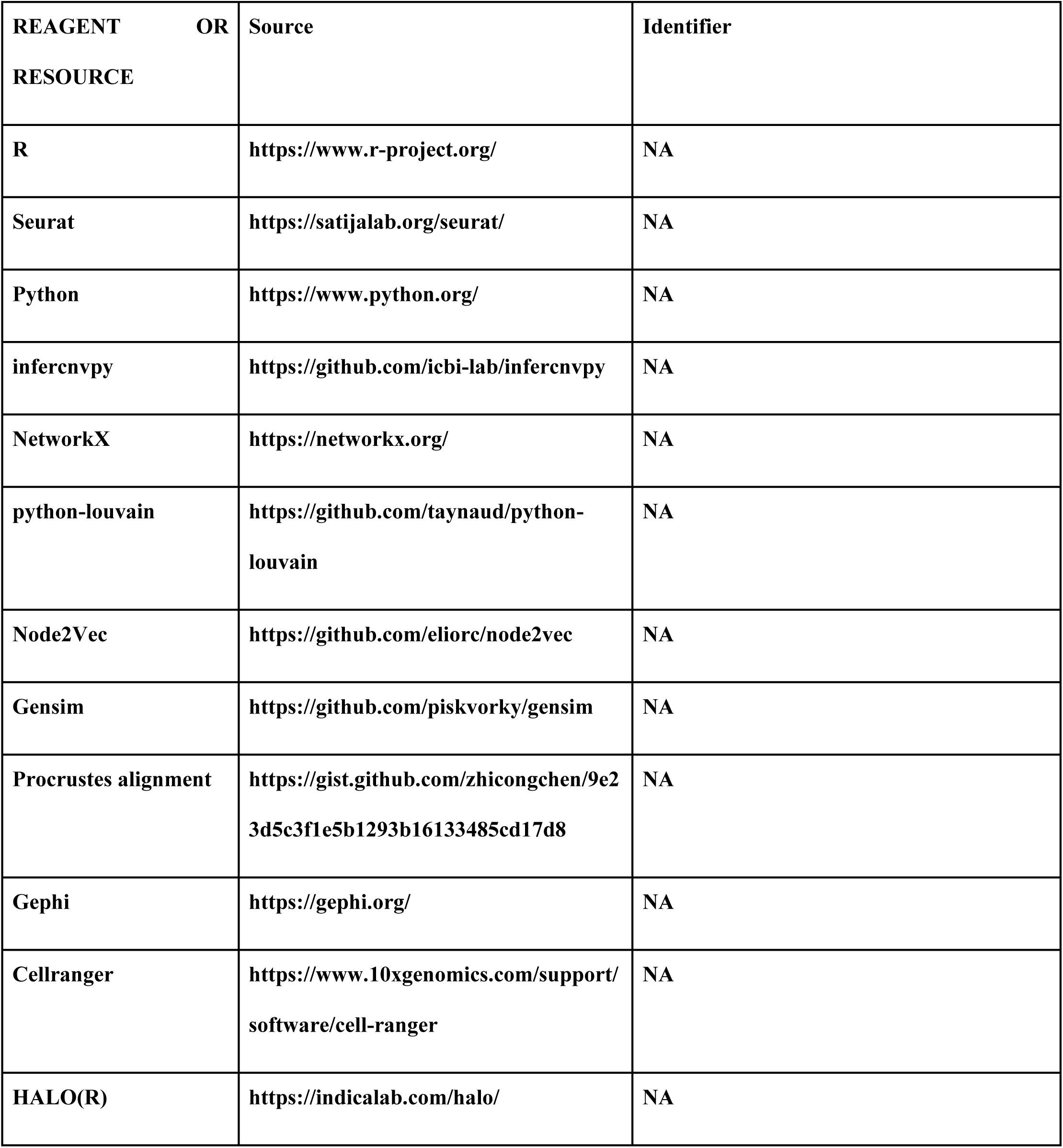

### Chemicals and drug compounds

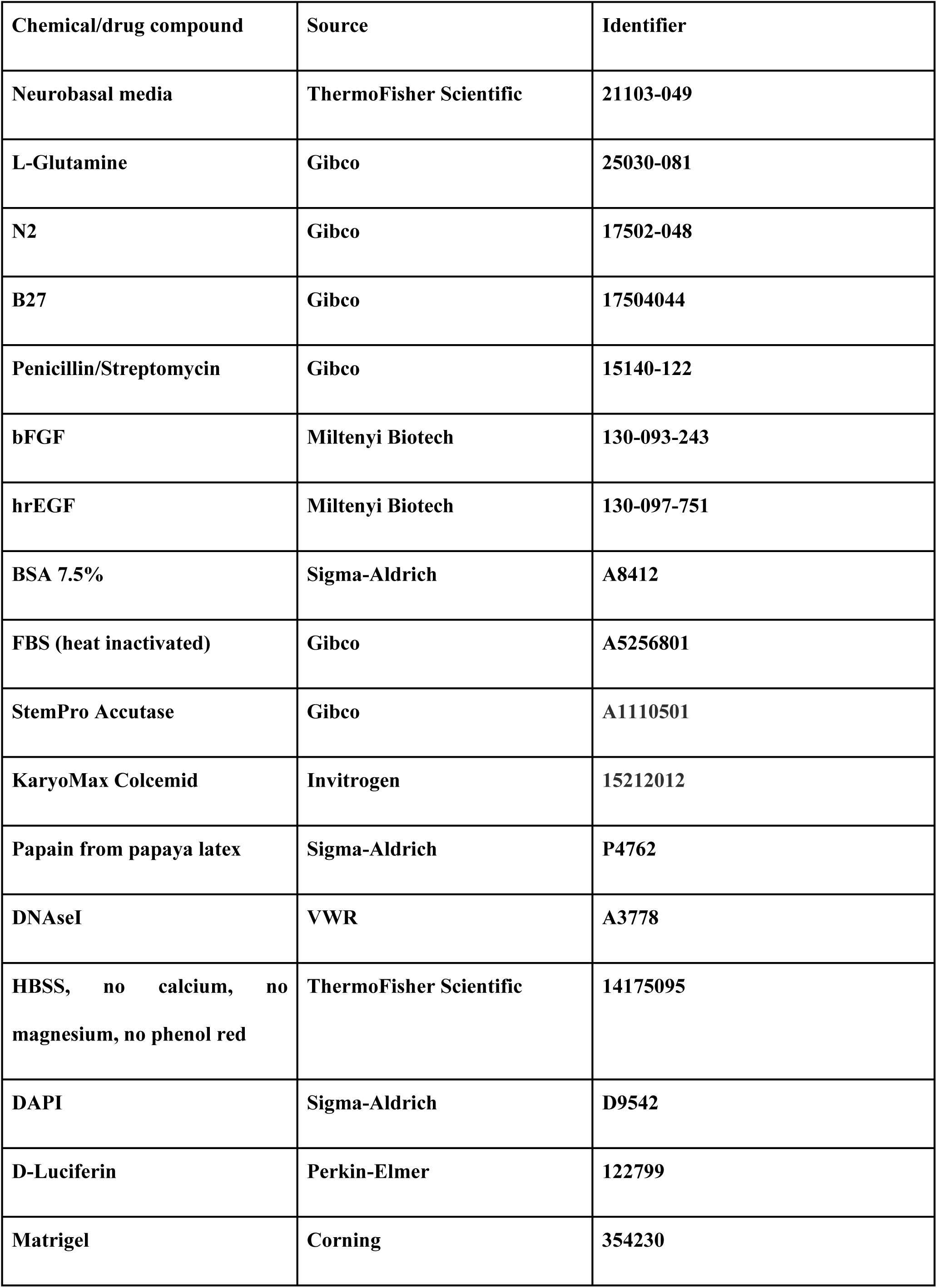
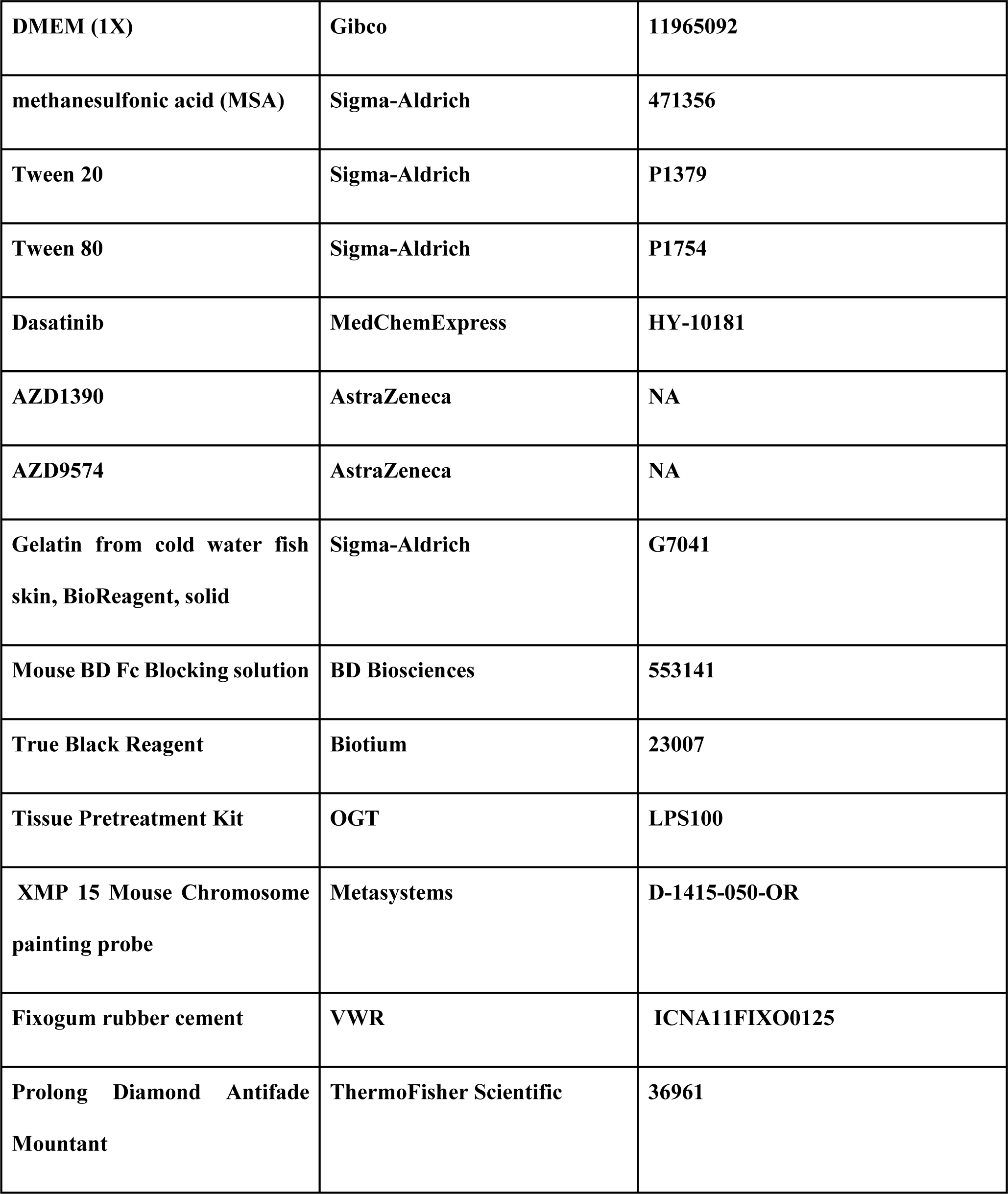

### Deposited data

|  |  |  |
| --- | --- | --- |
| **Raw single cell RNA-sequencing data** | **This paper** | **GEO: GSE252995** |

## REFERENCES

1. Santos-de-Frutos, K., and Djouder, N. (2021). When dormancy fuels tumour relapse. Communications Biology 2021 4:1 4, 1–12. 10.1038/s42003-021-02257-0.

2. Tyner, J.W., Haderk, F., Kumaraswamy, A., Baughn, L.B., Ness, B. Van, Liu, S., Marathe, H., Alumkal, J.J., Bivona, T.G., Chan, K.S., et al. Understanding Drug Sensitivity and Tackling Resistance in Cancer. 10.1158/0008-5472.CAN-21-3695.

3. Iragorri, N., Essue, B., Timmings, C., Keen, D., Bryant Md Phd, H., and Warren Md, G.W. (2020). The cost of failed first-line cancer treatment related to continued smoking in Canada. Current Oncology 27. 10.3747/co.27.5951.

4. Colleoni, M., Goldhirsch, A., Gianni, L., degli Infermi, O., Scientifico Romagnolo per lo Studio la Cura dei Tumori, I., Sun, Z., Price, K.N., Gelber, R.D., Foundation, R., Chan, H.T., et al. (2016). JOURNAL OF CLINICAL ONCOLOGY O R I G I N A L R E P O R T Annual Hazard Rates of Recurrence for Breast Cancer During 24 Years of Follow-Up: Results From the International Breast Cancer Study Group Trials I to V. 10.1200/JCO.2015.62.3504.

5. Marine, J.-C., Dawson, S.-J., and Dawson, M.A. (2020). Non-genetic mechanisms of therapeutic resistance in cancer. Nat Rev Cancer 20, 743–756. 10.1038/s41568-020-00302-4.

6. Salgia, R., and Kulkarni, P. (2018). The Genetic/Non-genetic Duality of Drug ‘Resistance’ in Cancer. Trends Cancer 4, 110–118. 10.1016/j.trecan.2018.01.001.

7. Holohan, C., Van Schaeybroeck, S., Longley, D.B., and Johnston, P.G. (2013). Cancer drug resistance: an evolving paradigm. Nature Reviews Cancer 2013 13:10 13, 714–726. 10.1038/nrc3599.

8. Stewart, A., Coker, E.A., Pölsterl, S., Georgiou, A., Minchom, A.R., Carreira, S., Cunningham, D., O’Brien, M.E.R., Raynaud, F.I., De Bono, J.S., et al. (2019). Differences in signaling patterns on PI3K inhibition reveal context specificity in KRAS mutant cancers. Mol Cancer Ther 18, 1396–1404. 10.1158/1535-7163.MCT-18-0727/358637/P/DIFFERENCES-IN-SIGNALING-PATTERNS-ON-PI3K.

9. Sun, X., Wang, S.C., Wei, Y., Yopp, A.C., Singal, A.G., and Correspondence, H.Z. (2017). Arid1a Has Context-Dependent Oncogenic and Tumor Suppressor Functions in Liver Cancer. Cancer Cell 32, 574–589. 10.1016/j.ccell.2017.10.007.

10. Jassim, A., Rahrmann, E.P., Simons, B.D., and Gilbertson, R.J. (2023). Cancers make their own luck: theories of cancer origins. Nature Reviews Cancer 2023, 1–15. 10.1038/s41568-023-00602-5.

11. Lopez, J.S., and Banerji, U. (2017). Combine and conquer: challenges for targeted therapy combinations in early phase trials. Nat. Rev. Clin. Oncol. 14, 57–66. 10.1038/nrclinonc.2016.96.

12. Al-Lazikani, B., Banerji, U., and Workman, P. (2012). Combinatorial drug therapy for cancer in the post-genomic era. Nat. Biotechnol. 30, 679–692. 10.1038/nbt.2284.

13. Jaaks, P., Coker, E.A., Vis, D.J., Edwards, O., Carpenter, E.F., Leto, S.M., Dwane, L., Sassi, F., Lightfoot, H., Barthorpe, S., et al. (2022). Effective drug combinations in breast, colon and pancreatic cancer cells. Nature 603, 166–173. 10.1038/s41586-022-04437-2.

14. Labrie, M., Brugge, J.S., Mills, G.B., and Zervantonakis, I.K. (2022). Therapy resistance: opportunities created by adaptive responses to targeted therapies in cancer. Nat Rev Cancer 22, 323–339. 10.1038/s41568-022-00454-5.

15. Pappo, A.S., Furman, W.L., Schultz, K.A., Ferrari, A., Helman, L., and Krailo, M.D. (2015). Rare tumors in children: Progress through collaboration. Journal of Clinical Oncology 33, 3047–3054. 10.1200/JCO.2014.59.3632.

16. Alvi, M.A., Wilson, R.H., and Salto-Tellez, M. (2017). Rare cancers: the greatest inequality in cancer research and oncology treatment. British Journal of Cancer 2017 117:9 117, 1255–1257. 10.1038/bjc.2017.321.

17. Halley, M.C., Stevens Smith, H., Ashley, E.A., Goldenberg, A.J., and Tabor, H.K. A call for an integrated approach to improve efficiency, equity and sustainability in rare disease research in the United States. 10.1038/s41588-022-01027-w.

18. Wright, K.D., Daryani, V.M., Turner, D.C., Onar-Thomas, A., Boulos, N., Orr, B.A., Gilbertson, R.J., Stewart, C.F., and Gajjar, A. (2015). Phase I study of 5-fluorouracil in children and young adults with recurrent ependymoma. Neuro Oncol 17, 1620–1627. 10.1093/neuonc/nov181.

19. Atkinson, J.M., Shelat, A.A., Carcaboso, A.M., Kranenburg, T.A., Arnold, L.A., Boulos, N., Wright, K., Johnson, R.A., Poppleton, H., Mohankumar, K.M., et al. (2011). An Integrated In Vitro and In Vivo High-Throughput Screen Identifies Treatment Leads for Ependymoma. Cancer Cell 20, 384–399. 10.1016/j.ccr.2011.08.013.

20. Gajjar, A., Stewart, C.F., Ellison, D.W., Kaste, S., Kun, L.E., Packer, R.J., Goldman, S., Chintagumpala, M., Wallace, D., Takebe, N., et al. (2013). Phase I study of vismodegib in children with recurrent or refractory medulloblastoma: A pediatric brain tumor consortium study. Clinical Cancer Research 19, 6305–6312. 10.1158/1078-0432.CCR-13-1425.

21. Robinson, G.W., Orr, B.A., Wu, G., Gururangan, S., Lin, T., Qaddoumi, I., Packer, R.J., Goldman, S., Prados, M.D., Desjardins, A., et al. (2015). Vismodegib exerts targeted efficacy against recurrent sonic hedgehog - Subgroup medulloblastoma: Results from phase II Pediatric Brain Tumor Consortium studies PBTC-025B and PBTC-032. Journal of Clinical Oncology 33, 2646–2654. 10.1200/JCO.2014.60.1591.

22. Romer, J.T., Kimura, H., Magdaleno, S., Sasai, K., Fuller, C., Baines, H., Connelly, M., Stewart, C.F., Gould, S., Rubin, L.L., et al. (2004). Suppression of the Shh pathway using a small molecule inhibitor eliminates medulloblastoma in Ptc1+/-p53-/- mice. Cancer Cell 6, 229–240. 10.1016/j.ccr.2004.08.019.

23. Rybinski, B., Hosgood, H.D., Wiener, S.L., and Weiser, D.A. (2020). Preclinical Metrics Correlate With Drug Activity in Phase II Trials of Targeted Therapies for Non-Small Cell Lung Cancer. Front Oncol 10, 2411. 10.3389/FONC.2020.587377/BIBTEX.

24. Kersten, K., de Visser, K.E., van Miltenburg, M.H., and Jonkers, J. (2017). Genetically engineered mouse models in oncology research and cancer medicine. EMBO Mol Med 9, 137–153. 10.15252/emmm.201606857.

25. Malbari, F. (2021). Pediatric Neuro-Oncology. Neurol Clin 39, 829–845. 10.1016/j.ncl.2021.04.005.

26. Cannon, D.M., Mohindra, P., Gondi, V., Kruser, T.J., and Kozak, K.R. (2015). Choroid plexus tumor epidemiology and outcomes: implications for surgical and radiotherapeutic management. J Neurooncol 121, 151–157. 10.1007/S11060-014-1616-X.

27. Takaoka, K., Cioffi, G., Waite, K.A., Finlay, J.L., Landi, D., Greppin, K., Kruchko, C., Ostrom, Q.T., and Barnholtz-Sloan, J.S. (2022). Incidence and survival of choroid plexus tumors in the United States. Neurooncol Pract, npac062. 10.1093/nop/npac062.

28. Zaky, W., and Finlay, J.L. (2018). Pediatric choroid plexus carcinoma: Biologically and clinically in need of new perspectives. Pediatr Blood Cancer 65, e27031. 10.1002/pbc.27031.

29. Barlow-Krelina, E., Chen, Y., Yasui, Y., Till, C., Gibson, T.M., Ness, K.K., Leisenring, W.M., Howell, R.M., Nathan, P.C., Oeffinger, K.C., et al. (2020). Consistent Physical Activity and Future Neurocognitive Problems in Adult Survivors of Childhood Cancers: A Report From the Childhood Cancer Survivor Study. Journal of Clinical Oncology 38, 2041–2052. 10.1200/JCO.19.02677.

30. Tong, Y., Merino, D., Nimmervoll, B., Gupta, K., Wang, Y.D., Finkelstein, D., Dalton, J., Ellison, D.W., Ma, X., Zhang, J., et al. (2015). Cross-Species Genomics Identifies TAF12, NFYC, and RAD54L as Choroid Plexus Carcinoma Oncogenes. Cancer Cell 27, 712–727. 10.1016/j.ccell.2015.04.005.

31. Nimmervoll, B. V, Boulos, N., Bianski, B., Dapper, J., DeCuypere, M., Shelat, A., Terranova, S., Terhune, H.E., Gajjar, A., Patel, Y.T., et al. (2018). Establishing a Preclinical Multidisciplinary Board for Brain Tumors. Clinical Cancer Research 24, 1654–1666. 10.1158/1078-0432.CCR-17-2168.

32. Dani, N., Herbst, R.H., McCabe, C., Green, G.S., Kaiser, K., Head, J.P., Cui, J., Shipley, F.B., Jang, A., Dionne, D., et al. (2021). A cellular and spatial map of the choroid plexus across brain ventricles and ages. Cell 184, 3056–3074.e21. 10.1016/J.CELL.2021.04.003.

33. Johnson, R.A., Wright, K.D., Poppleton, H., Mohankumar, K.M., Finkelstein, D., Pounds, S.B., Rand, V., Leary, S.E.S., White, E., Eden, C., et al. (2010). Cross-species genomics matches driver mutations and cell compartments to model ependymoma. Nature 466, 632–636. 10.1038/nature09173.

34. Gibson, P., Tong, Y., Robinson, G., Thompson, M.C., Currle, D.S., Eden, C., Kranenburg, T.A., Hogg, T., Poppleton, H., Martin, J., et al. (2010). Subtypes of medulloblastoma have distinct developmental origins. Nature 468, 1095–1099. 10.1038/nature09587.

35. Behjati, S., Gilbertson, R.J., and Pfister, S.M. (2021). Maturation Block in Childhood Cancer. Cancer Discov 11, 542–544. 10.1158/2159-8290.cd-20-0926.

36. Jessa, S., Blanchet-Cohen, A., Krug, B., Vladoiu, M., Coutelier, M., Faury, D., Poreau, B., De Jay, N., Hébert, S., Monlong, J., et al. (2019). Stalled developmental programs at the root of pediatric brain tumors. Nat Genet 51, 1702–1713. 10.1038/s41588-019-0531-7.

37. Flanary, V.L., Fisher, J.L., Wilk, E.J., Howton, T.C., and Lasseigne, B.N. (2023). Computational Advancements in Cancer Combination Therapy Prediction. 10.1200/PO.23.00261. 10.1200/PO.23.00261.

38. Grover, A., and Leskovec, J. (2016). node2vec: Scalable Feature Learning for Networks. Kdd 2016, 855–864. 10.1145/2939672.2939754.

39. Mikolov, T., Chen, K., Corrado, G., and Dean, J. (2013). Efficient Estimation of Word Representations in Vector Space. 1st International Conference on Learning Representations, ICLR 2013 - Workshop Track Proceedings.

40. Nowakowski, T.J., Bhaduri, A., Pollen, A.A., Alvarado, B., Mostajo-Radji, M.A., Di Lullo, E., Haeussler, M., Sandoval-Espinosa, C., Liu, S.J., Velmeshev, D., et al. (2017). Spatiotemporal gene expression trajectories reveal developmental hierarchies of the human cortex. Science 358, 1318–1323. 10.1126/SCIENCE.AAP8809.

41. A Translational Repression Complex in Developing Mammalian Neural Stem Cells that Regulates Neuronal Specification 10.1016/j.neuron.2017.12.045.

42. Schmitz, K.M., Schmitt, N., Hoffmann-Rohrer, U., Schäfer, A., Grummt, I., and Mayer, C. (2009). TAF12 Recruits Gadd45a and the Nucleotide Excision Repair Complex to the Promoter of rRNA Genes Leading to Active DNA Demethylation. Mol Cell 33, 344–353. 10.1016/J.MOLCEL.2009.01.015.

43. Swagemakers, S.M.A., Essers, J., de Wit, J., Hoeijmakers, J.H.J., and Kanaar, R. (1998). The Human Rad54 Recombinational DNA Repair Protein Is a Double-stranded DNA-dependent ATPase*. Journal of Biological Chemistry 273, 28292–28297. 10.1074/jbc.273.43.28292.

44. Mohankumar, K.M., Currle, D.S., White, E., Boulos, N., Dapper, J., Eden, C., Nimmervoll, B., Thiruvenkatam, R., Connelly, M., Kranenburg, T.A., et al. (2015). An in vivo screen identifies ependymoma oncogenes and tumor-suppressor genes. Nat Genet 47, 878–887. 10.1038/ng.3323.

45. Durant, S.T., Zheng, L., Wang, Y., Chen, K., Zhang, L., Zhang, T., Yang, Z., Riches, L., Trinidad, A.G., Fok, J.H.L., et al. (2018). The brain-penetrant clinical ATM inhibitor AZD1390 radiosensitizes and improves survival of preclinical brain tumor models. Sci Adv 4. 10.1126/SCIADV.AAT1719.

46. Jucaite, A., Stenkrona, P., Cselényi, Z., De Vita, S., Buil-Bruna, N., Varnäs, K., Savage, A., Varrone, A., Johnström, P., Schou, M., et al. (2021). Brain exposure of the ATM inhibitor AZD1390 in humans—a positron emission tomography study. Neuro Oncol 23, 687–696. 10.1093/neuonc/noaa238.

47. Saha, S., Jungas, T.T., Ohayon, D., Audouard, C., Ye, T., Fawal, M.A., and Davy, A. (2023). Dihydrofolate reductase activity controls neurogenic transitions in the developing neocortex. Development 150. 10.1242/DEV.201696.

48. Gangoso, E., Southgate, B., Bradley, L., Rus, S., Galvez-Cancino, F., McGivern, N., Güç, E., Kapourani, C.-A., Byron, A., Ferguson, K.M., et al. (2021). Glioblastomas acquire myeloid-affiliated transcriptional programs via epigenetic immunoediting to elicit immune evasion. Cell 184, 2454–2470.e26. 10.1016/j.cell.2021.03.023.

49. Kirschenbaum, D., Xie, K., Weiss, T., and Weiner, A. Article Time-resolved single-cell transcriptomics defines immune trajectories in glioblastoma In brief. 10.1016/j.cell.2023.11.032.

50. Duggan, S.P., Garry, C., Behan, F.M., Phipps, S., Kudo, H., Kirca, M., Zaheer, A., McGarrigle, S., Reynolds, J. V., Goldin, R., et al. (2018). siRNA Library Screening Identifies a Druggable Immune-Signature Driving Esophageal Adenocarcinoma Cell Growth. Cell Mol Gastroenterol Hepatol 5, 569–590. 10.1016/J.JCMGH.2018.01.012.

51. Deczkowska, A., Weiner, A., and Amit, I. (2020). The Physiology, Pathology, and Potential Therapeutic Applications of the TREM2 Signaling Pathway. Cell 181, 1207–1217. 10.1016/J.CELL.2020.05.003.

52. Menche, J., Sharma, A., Kitsak, M., Ghiassian, S.D., Vidal, M., Loscalzo, J., and Barabási, A.L. (2015). Disease networks. Uncovering disease-disease relationships through the incomplete interactome. Science 347, 841. 10.1126/SCIENCE.1257601.

53. Cheng, F., Kovács, I.A., and Barabási, A.L. (2019). Network-based prediction of drug combinations. Nat Commun 10. 10.1038/S41467-019-09186-X.

54. Federico, A., Fratello, M., Scala, G., Möbus, L., Pavel, A., Del Giudice, G., Ceccarelli, M., Costa, V., Ciccodicola, A., Fortino, V., et al. (2022). Integrated Network Pharmacology Approach for Drug Combination Discovery: A Multi-Cancer Case Study. Cancers (Basel) 14. 10.3390/CANCERS14082043.

55. Amzallag, A., Ramaswamy, S., and Benes, C.H. (2019). Statistical assessment and visualization of synergies for large-scale sparse drug combination datasets. BMC Bioinformatics 20. 10.1186/S12859-019-2642-7.

56. Celebi, R., Bear Don’t Walk, O., Movva, R., Alpsoy, S., and Dumontier, M. (2019). In-silico Prediction of Synergistic Anti-Cancer Drug Combinations Using Multi-omics Data. Sci Rep 9. 10.1038/S41598-019-45236-6.

57. Mantovani, A., Marchesi, F., Malesci, A., Laghi, L., and Allavena, P. (2017). Tumour-associated macrophages as treatment targets in oncology. Nat Rev Clin Oncol 14, 399–416. 10.1038/nrclinonc.2016.217.

58. Ulland, T.K., Song, W.M., Huang, S.C.C., Ulrich, J.D., Sergushichev, A., Beatty, W.L., Loboda, A.A., Zhou, Y., Cairns, N.J., Kambal, A., et al. (2017). TREM2 Maintains Microglial Metabolic Fitness in Alzheimer’s Disease. Cell 170, 649–663.e13. 10.1016/J.CELL.2017.07.023.

59. Mohankumar, K.M., Currle, D.S., White, E., Boulos, N., Dapper, J., Eden, C., Nimmervoll, B., Thiruvenkatam, R., Connelly, M., Kranenburg, T.A., et al. (2015). An in vivo screen identifies ependymoma oncogenes and tumor-suppressor genes. Nat Genet 47, 878–887. 10.1038/ng.3323.

60. Zhang, C., Cerveira, E., Rens, W., Yang, F., and Lee, C. (2018). Multicolor Fluorescence In Situ Hybridization (FISH) Approaches for Simultaneous Analysis of the Entire Human Genome. Curr Protoc Hum Genet 99, e70. 10.1002/CPHG.70.

61. Andreatta, M., and Carmona, S.J. (2021). UCell: Robust and scalable single-cell gene signature scoring. Comput Struct Biotechnol J 19, 3796–3798. 10.1016/j.csbj.2021.06.043.

62. Schönemann, P.H. (1966). A generalized solution of the orthogonal procrustes problem. Psychometrika 31, 1–10. 10.1007/BF02289451/METRICS.

